# Longer projected droughts will impair the recovery of tropical seedlings and their leaf microbiota

**DOI:** 10.64898/2026.08.26.746266

**Authors:** Marion Boisseaux, Jean-Yves Goret, Benoit Burban, Valérie Troispoux, Alice Bordes, Jocelyn Cazal, Saint-Omer Cazal, Sabrina Coste, Clément Stahl, Heidy Schimann

## Abstract

The increasingly severe droughts in the Amazon Basin make it urgent to understand the resilience of tropical tree species and their microbiota. Plant-associated fungi and bacteria (i.e. *extended phenotype)* modulate drought stress for their hosts, but their role in recovery dynamics remains poorly understood.

To test the impact of different drought durations on the recovery of both physiology and microbiota of tropical trees, we followed the responses of nearly 1,000 seedlings belonging to seven tropical tree species of seasonally flooded (SF) forests in a greenhouse experiment. Seedlings were subjected to different droughts, reflecting a current, a projected and an extreme drought scenario of the French Guiana climate. Plant responses were monitored after the drought and after rewetting. Plant performance was estimated through leaf gas exchange, photochemical functioning, leaf water potentials and water-related traits as well as morphological traits. Bacterial and fungal leaf communities were characterized with respectively 16S and ITS2 markers using high-throughput sequencing.

Increasing the duration of the drought reduced the ability of plants to recover physiological functions, with differences among species which were only partially predicted by their drought tolerance strategies. Bacterial diversity increased in most plant host species after mild drought but not under the most severe stress. Bacterial dispersion and turnover responses were strongly host species-specific, without a general directional pattern across species. Fungal communities showed greater compositional stability, but exhibited consistently higher turnover compared to bacterial communities during both drought and recovery, with no convergence toward control composition. Finally, none of the recovery networks mirrored the architecture of the control network, regardless of prior drought duration, demonstrating that the integrated extended phenotype does not recover even when individual traits show signs of recovery. Our results reveal that both physiological recovery and microbial community recovery are strongly shaped by the plant host species identity and drought duration

This study widens knowledge of SF tropical forests, vulnerable habitats in the context of climate change, through the lens of the associated microbial communities and functional traits. Beyond the effects of an increasingly uncertain climate combined with a rise in the frequency of extreme events, our study places emphasis on including tree species extended phenotypes in considering their recovery dynamics.

**Graphical Abstract:** 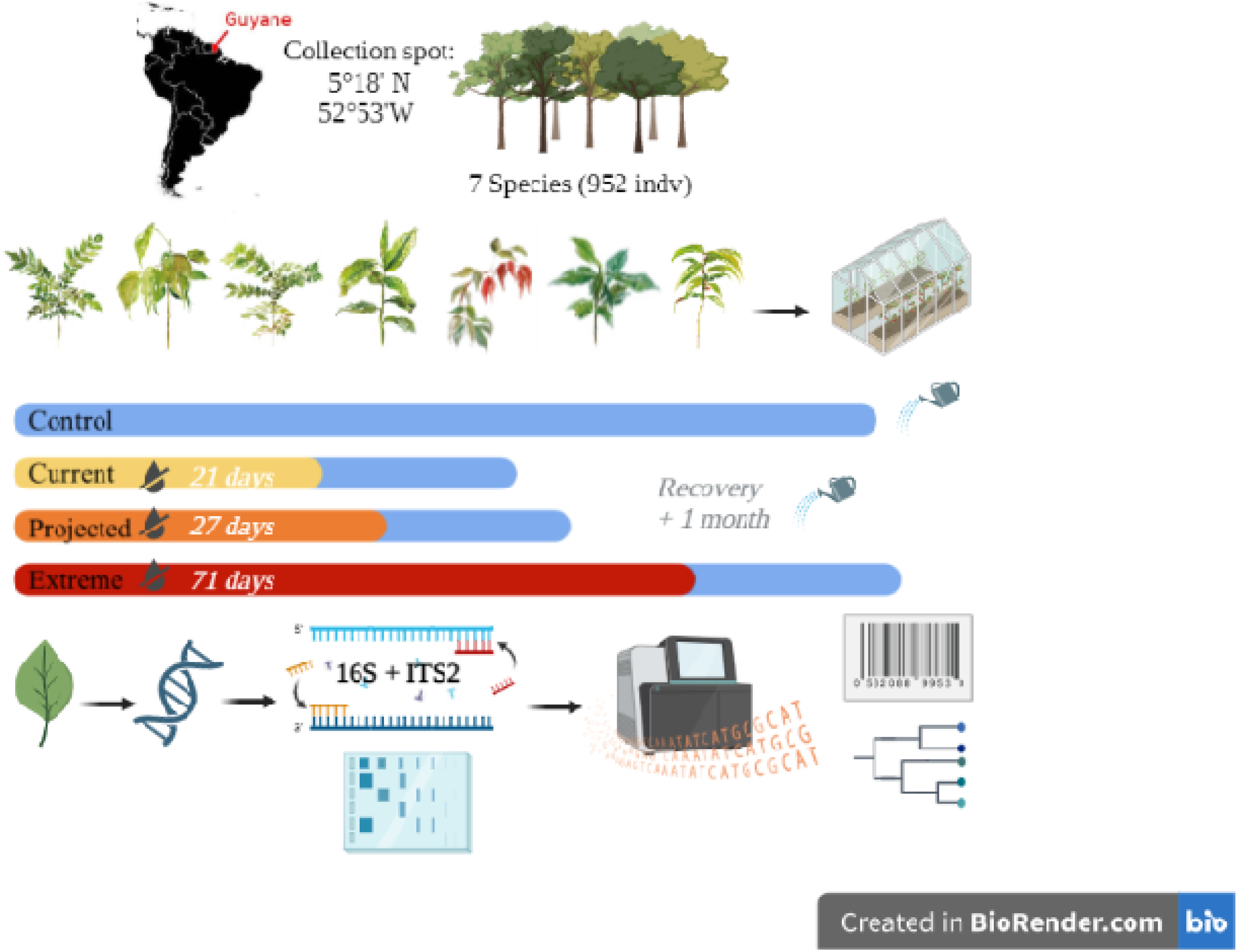

## Introduction

Climate change challenges the ability of ecosystems to recover from disturbances (Phillipot *et al*., 2021). The global increase of drought events has become one of the main threats to tropical forests (Calvin *et al*., 2023), impacting forest productivity and tree mortality (Phillips *et al*., 2010; Brodribb *et al*., 2020). While previous studies have largely focused on how trees cope with drought (O’Grady *et al*., 2013; Delzon 2015; Brodribb, 2016; Volaire 2018; Blackman *et al*., 2019; Ziegler *et al*., 2023; Volaire *et al*., 2026), predicting the long-term resilience of tropical forests also requires a mechanistic understanding of post-drought recovery mechanisms (Yin and Bauerle 2017).

In trees, drought triggers major physiological and structural responses which progressively constrain plant water and carbon functioning (Bonal *et al*., 2016; McDowell 2008). Droughts limit carbon acquisition as stomatal conductance (*g*_s_) and carbon assimilation (*A*_sat_) decrease, consequently lowering the absolute growth rate (AGR) (Delzon, 2015; Volaire, 2018). These responses under short droughts are generally reversible upon rewatering (Chen *et al*., 2010; Manzi *et al*., 2022). Photosynthetic capacity, evaluated by the maximum efficiency of photosystem II (*F*_v_/*F*_m_), often declines under moderate drought but can recover gradually upon rewatering as biochemical components are rebuilt, but recovery trajectories can vary among species (Kirschbaum 1988; Gallé *et al*., 2007; Manzi *et al*., 2022). In contrast, longer droughts resulting in more severe water stress can result in structural damage, causing recovery to be slow. Prolonged water deficit disrupts the plant water status and reduces the leaf water storage capacity, as reflected by declines in the leaf water potential and relative water content (RWC). RWC is a direct measurement of the leaf water status (Sapes *et al*., 2019) and could be an indicator of the tree’s capacity to recover (Mantova *et al*., 2021). As dehydration lasts longer, hydraulic functions become highly constrained, such that the rate of leaf hydraulic repair ultimately limits the post-drought recovery of gas exchange (Blackman *et al*., 2009). If drought becomes too severe, xylem embolism can occur, provoking irreversible loss of conductivity (Brodribb and Cochard 2009). Thus, the reversibility of plant functions (decrease in carbon assimilation, stomatal closure, down-regulation of photosynthetic capacity, reduction of leaf water potential) will depend on drought duration.

Drought recovery not only depends on the tree species physiology but also their extended phenotype; *i.e* their associated microbial communities, which play a vital role in plant health (Vandenkoornhuyse *et al*., 2015; Santos-Medellín *et al*., 2017; Naylor *et al*., 2017; Trivedi *et al*., 2020). Endophytes *sensus* Hardoim *et al*. (2015), dominated by bacteria and fungi (including yeasts) (Rosado *et al*., 2018; Stone *et al*., 2018; Chaudhry *et al*., 2021), and have been involved in plant responses via diverse mechanisms, including growth promotion by hormones, modulation of the plant immune system, and enhancement of biotic and abiotic stress tolerance (Zhang *et al*., 2021). Although the majority of work in plant-microbiota interactions has focused on the root compartment (Naylor *et al*., 2017; Santos-Medellin *et al*., 2017; Xu *et al*., 2018; Li *et al*., 2025; Wu *et al*., 2026), the leaf *i.e.* the phyllosphere, is also expected to play a key role in plant recovery capacity. The phyllosphere environment is subject to more fluctuating environmental conditions such as light, temperature and relative humidity compared to the soil or rhizosphere (Vorholt, 2012). Drought can alter the diversity of leaf microbiota (Bechtold *et al*., 2021; Karasov *et al*., 2024), including shifts in both bacterial and fungal communities. Since bacteria and fungi differ in their ecological strategies, studies have shown that bacteria are generally more sensitive to drought stress than fungi (Naylor and Colemann-Derr, 2018; Yang *et al*., 2026). Drought was found to reduce leaf bacterial richness and only the relative abundance of dominant fungal taxa (Debray *et al*., 2022). In grass roots, drought has been shown to reduce bacterial diversity, with communities becoming dominated by Actinobacteria and other stress-tolerant lineages (Xu *et al*., 2018; Santos-Medellín *et al*., 2017). Beyond diversity, drought strongly affects the dispersion of communities (*i.e.* variation among plant hosts) and their turnover (*i.e.* replacement of taxa), two complementary metrics that capture distinct aspects of the microbiota dynamics during disturbance and recovery (Jurburg *et al*., 2024). Drought is expected to reduce dispersion of microorganisms by selecting for stress-tolerant taxa, thereby homogenising communities across host plants (Debray *et al*., 2022). Following this drought induced-decrease, dispersion is expected to increase during the recovery phase as microbial communities re-diversify (Jurburg *et al*., 2024). However, after a certain stress threshold, the plant loses control over its microbiota, leading to dysbiosis with an increase of dispersion under more severe droughts (Arnault *et al*., 2023), and a decrease during recovery. Concerning turnover, drought-tolerant but potentially functionally distinct taxa replace previous microbial taxa, resulting in the loss of core microbial members and a shift of microbial composition which increases turnover (Santos-Medellin *et al*., 2017; Jurburg *et al*., 2024; Yang *et al*., 2026). Following rewatering, microbial communities are expected to converge back to their pre-drought composition upon rewatering (*i.e.* decreasing turnover). However, the extent of this compositional recovery is expected to decrease with increasing drought duration, as more prolonged droughts lead to more irreversible differences (in roots: Santos-Medellin *et al*., 2021; in leaves: Bechtold *et al*., 2021).

Recovery strategies are usually defined as a return to initial states (measured against a control treatment or pre-stress conditions), or reaching a new stable-functioning state (Ingrisch and Bahn 2018; Van Meerbeek *et al*., 2021; Dakos and Kefi 2022). Recovery strategies fall into four categories: no recovery, partial, complete or over-compensation recovery (Ruehr *et al*., 2019). Since drought duration can determine the reversibility of plant physiological functions, we predicted that (i) increasing drought duration would progressively constrain the recovery of water- and carbon-related functions, leading to a complete recovery after short and moderate droughts, but incomplete or no recovery after a longer drought (Figure 1A). We also expect species identities to drive differences in recovery responses. For microbial recovery, we predicted that (ii) microbial diversity would increase back to pre-drought levels upon rewatering. We also predicted that (iii) dispersion should respond in the opposite direction of drought effects, either increase toward control levels after a drought-induced reduction, or decrease after dysbiosis (Figure 1B). Finally, we expected (iv) microbial communities to show decreasing turnover (*i.e.*, convergence back toward control composition) during recovery and that the extent of this recovery should decrease with increasing drought duration (Figure 1C).

**Figure 1:**
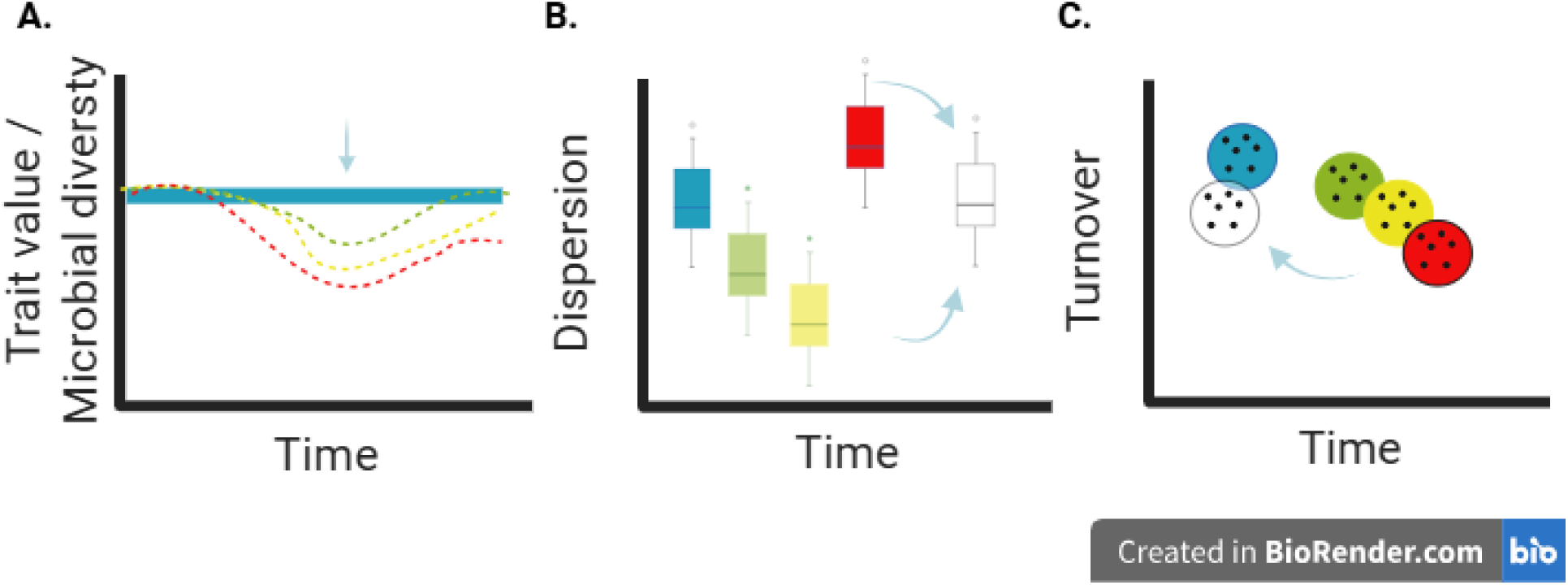
Hypotheses concerning plant trait and microbial community drought recovery dynamics. Line and boxplot colors indicate control (blue), increasing drought duration (green, yellow, red), and recovery (white). Arrows indicate recovery. **A: Trait variation or microbial diversity patterns.** We hypothesized that the capacity of water- and carbon-related traits to return to control values would decrease with increasing drought duration. Microbial diversity would decrease with increasing drought duration, followed by a return toward control levels after rewatering. **B: Dispersion patterns**. Dispersion represents the variability among replicates within each treatment. We hypothesized that drought would reduce dispersion as strong environmental filtering selects for stress-tolerant taxa, reducing community heterogeneity, until a certain threshold (dysbiosis). Upon recovery, dispersion should return towards control levels. **C: Turnover patterns.** Turnover represents how far recovering communities are from the control. We expected microbial communities to show decreasing turnover values, approaching those of the control communities during recovery. The extent of this recovery should decrease with increasing drought duration.

To test our hypotheses, we conducted an original greenhouse experiment of nearly 1,000 seedlings belonging to seven tropical tree species of seasonally flooded forest (SF). SF forests represent almost 50% of Amazon basin area (Costa *et al*., 2023) a widespread but understudied tropical forest type, crucial to predicting the resilience of Amazonian forest to climate change. Seedlings were subjected to drought treatments that reflected current, projected and extreme drought scenarios of the climate in French Guiana. Each drought scenario was followed by a recovery period of one month. We monitored both the physiological responses of seedlings as well as leaf fungal and bacteria communities. Our analyses were structured along three recovery dimensions: the recovery of (i) leaf trait values, (ii) microbial community diversity and (iii) composition through dispersion and turnover metrics. Finally, in an effort to explore the response of the extended phenotype, we used a network approach which jointly captures how the physiology of the plant and its endophytes are structured and co-re-organise during and after drought.

## Materials and methods

### Sampling and growing conditions

Seedlings were collected at the Paracou field station (5°18′N, 52°53′W) located in French Guiana, between January and September of 2021. This site is characterized by an average annual rainfall of 2875 mm and a mean air temperature of 26°C (Gourlet-Fleury *et al*., 2004). Seasonally flooded forests (SF) are located in the bottom lands and are characterized by relatively fertile and sandy soils. The water table is never observed to descend below 60 cm depth, remaining at the soil surface for at least two consecutive months each year during the rainy season (Ferry *et al*., 2010). More than 2,000 1-year-old seedlings were sampled in SF forest habitats exclusively belonging to the following species: *Eperua falcata* Aubl., *Iryanthera hostmannii* (Benth.) Warb., *Jacaranda copaia subsp. copaia* Aubl., *Pterocarpus officinalis* Jacq., *Symphonia globulifera* L.f., *Tachigali melinonii* (Harms) Zarucchi & Herend, *Virola surinamensis* (Rol. ex Rottb.) Warb. (Fougnies *et al*., 2007; Baraloto *et al*., 2007; Baraloto *et al*., 2021) (Figure S1, Table 1). Seedlings were pooled in batches, organized by species and GPS position (Each unique GPS position corresponded to 1 mother tree) (Figure S2, Table S1). After collecting the seedlings, they were immediately brought back to the campus for grounding at the ECOFOG research facilities in Kourou. Seedlings were then directly transplanted into 4 L pots containing a 2:1 (v/v) mixture of brown forest ferralitic clay soil and sand. Soil proportions were similar between the forest soil (83.2% sand; 79.7% clay; 7.1% limon) and the greenhouse soil (85.3% sand; 7% clay and 7.7% limon). More detailed soil analyses can be found in Table S2. Seedlings were then grown in a shadehouse under ca. 7% of total irradiance for at least four months and irrigated two times a day (8 am and 6 pm during 5 min). A total of 952 seedlings were then placed into the greenhouse for the experiment.

**Table 1:** Focal species of the study (Baraloto et al., 2021).

| Family | Genus | Species | Habitat preference |
| --- | --- | --- | --- |
| FABACEAE | *Eperua* | *falcata* | SF forests |
| MYRISTICACEAE | *Iryanthera* | *hostmannii* | SF forests |
| BIGNONIACEAE | *Jacaranda* | *copaia* | generalist |
| FABACEAE | *Pterocarpus* | *officinalis* | SF forests |
| CLUSIACEAE | *Symphonia* | *globulifera* | SF forests |
| FABACEAE | *Tachigali* | *melinonii* | generalist |
| MYRISTICACEAE | *Virola* | *surinamensis* | SF forests |

### Experimental design

To investigate how different drought durations affect recovery capacity in SF seedlings, we exposed SF seedlings to three controlled dry-down events, followed by a recovery period of one month until soil volumetric water content was back to normal conditions (*i.e.* field capacity) (Figure S5).

Historical rainfall data recorded at Sinnamary Weather Station (MétéoFrance) over the past 64 years (1955-2019), enabled us to define realistic drought durations. The annual mean ± SD maximum number of consecutive days without rainfall (detection limit 0.2 mm) in a dry season (August-November) was 21 ± 5 days, and climate projections forecast a 30 % reduction in precipitation in the Amazon region by 2100 (Calvin *et al*., 2023), corresponding to 27 consecutive days without rainfall. As an extreme scenario, we used the maximum number of 71 consecutive days without rainfall recorded in 1976 (Figure S4).

1. Well-watered every 2-3 days to field capacity (C0).
2. 21 days without water + recovery period of one month (D1 to R1).
3. 27 days without water + recovery period of one month (D2 to R2).
4. 71 days without water + recovery period of one month (D3 to R3).

In total, 952 seedlings were used for the experiment with 136 individual seedlings per species, in a completely randomized block design (see Figure S6 for further details). Individuals for each species were randomly assigned within each block unit, in order to mix mother tree origin within treatments and avoid greenhouse seedling position effects.

Light, temperature and air relative humidity in the greenhouse were monitored using Environmental HOBO sensors (model UA-002-64, HOBO Pendant Tem Light—64 k and model U23–001, HOBO Pro V2 Temp/RH Datalogger, Amanvillers, France). The mean diurnal relative humidity was 82.9%, ranging from 60.5% to 95.3%. The mean diurnal temperature was 33.4°C, ranging from 24.7°C to 42.6°C. The light intensity was 33.8% of full external irradiance.

### Soil analyses

As seedlings were directly transplanted into 4 L pots filled with a mixture of brown forest ferralitic clay soil and sand, soil samples from several pots were compared to *in natura* forest soil samples where the seedlings were taken at the Paracou field station (Table S2; Figure S3). The soil analyses were carried out by the Soil Analyses Laboratory of the Cirad, Montpellier. Overall, the substrate used in our experiment is slightly less acidic than *in natura* conditions.

### Functional traits

Functional traits were measured at key sampling dates (Table S3) on five randomly chosen plants per treatment and three control plants per species, except for basal diameter (D, mm) and height (H, cm), which were measured in all individuals at each campaign. A total of 16 functional traits were quantified (Table 2).

**Table 2:** Traits used in this study and their functional mechanisms.

| Trait | Abbreviation | Unit | Type | Mechanisms |
| --- | --- | --- | --- | --- |
| Stomatal conductance | $g_s$ | $\text{mmol} \cdot \text{m}^{-2} \cdot \text{s}^{-1}$ | Physiological | Carbon uptake & water status |
| Carbon assimilation rate under saturated light | $A_{\text{sat}}$ | $\mu\text{mol CO}_2 \cdot \text{m}^{-2} \cdot \text{s}^{-1}$ | Physiological | Carbon uptake |
| Evapotranspiration | E | $\text{mmol H}_2\text{O} \cdot \text{m}^{-2} \cdot \text{s}^{-1}$ | Physiological | Water loss |
| Water use efficiency | WUE | $\mu\text{mol CO}_2 \cdot \text{mmol}^{-1} \text{H}_2\text{O}$ | Physiological | Carbon uptake and water loss |
| Maximum quantum efficiency of photosystem II photochemistry | $F_v/F_m$ | unitless | Physiological | Photosynthetic activity |
| Chlorophyll content | Chl | SPAD unit | Physiological | Photosynthetic activity |
| Relative water content | RWC | % | Physiological | Water capacitance |
| Midday leaf water potential | $\Psi_{\text{midday}}$ | MPa | Physiological | Water status and flow |
| Total leaf area | $LA_T$ | $\text{cm}^2$ | Morphological | Resource capture |
| Leaf area | LA | $\text{cm}^2$ | Morphological | Resource capture |
| Specific leaf area | SLA | $\text{cm}^2 \cdot \text{g}^{-1}$ | Morphological | Resource capture |
| Leaf thickness | LT | mm | Morphological | Water capacitance |
| Root to shoot ratio | R:S | unitless | Morphological | Resource storage |
| Diameter | D | mm | Morphological | Resource storage, mechanical support |
| Height | H | cm | Morphological | Resource capture, mechanical support |
| Absolute growth rate | AGR | $\text{mm} \cdot \text{d}^{-1}$ | Morphological | Resource storage, mechanical support |

The absolute growth rate (mm.d^-1^) was calculated using basal diameter measurements, after the drought phase as *equation 1* and after the recovery phase as *equation 2*.

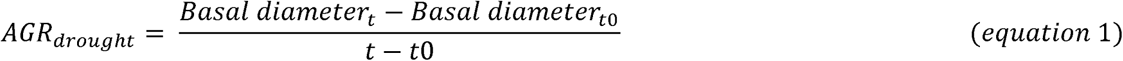

In equation 1, *t* corresponds to 21 days (D1), 27 days (D2) and 71 days (D3) after drought onset, and *t0* when water was withheld.

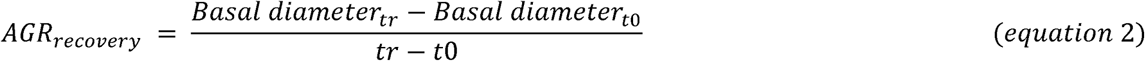

In equation 2, *t* corresponds to 51 days (recovery from D1, R1), 57 days (recovery from D2, R2) and 101 days (recovery from D3, R3).

### Gas exchange and chlorophyll fluorescence measurements

Stomatal conductance (*g*_s_, mmol.m^−2^.s^−1^) was measured between 8:00 a.m. and 11:00 a.m. using a porometer (AP4, Delta-T Devices, Cambridge, UK) on one leaf per plant (control and treatment), under ambient light and temperature conditions. The carbon assimilation rate under saturated light (*A*_sat_, µmol CO_2_. m^-2^.s^-1^), evapotranspiration (E, mmol H_2_O.m^-2^.s^-1^) and water use efficiency (WUE, µmol CO_2_. mmol^-1^ H_2_O) were performed with a Portable Photosynthesis System (CIRAS-3, PP-system, Amesbury, MA, USA). The parameters used were 1000 µmol.photons.m^-2^.s^-1^ PAR, 420 ppm CO_2_ concentration (actual ambient CO_2_ concentration measured on 21/07/2020 at the Paracou Flux Tower) and 28°C for the leaf chamber block temperature. The maximum photochemical efficiency of photosystem II (*F_v_/F_m_*, unitless) was measured using a portable pulse-modulated fluorometer (Mini-PAM II, WALZ, Effeltrich, Germany). Leaves were dark adapted for 30 min before measurements using leaf clips. Leaf chlorophyll content (Chl, SPAD unit) was determined with a SPAD-502 instrument.

### Water-related traits

Midday leaf water potential (Ψ_midday_, MPa) was measured between 11:00 a.m. and 1:00 p.m. using leaf-cutter thermocouple psychrometers (76–1.5VC, JRD Merill, USA) connected to a Psypro water potential data logger (Psypro, Wescor Inc., Logan, USA). Three leaf discs (diameter 5 mm) were sampled on one leaf seedling, sealed in the chamber, equilibrated at 25°C for at least 5 hours in water baths. Leaf water potential was then calculated from the initial slope of the psychometric response curve and previous calibrations with NaCl solutions.

We measured leaf relative water content (RWC, %) for which we obtained saturated weights by rehydrating the leaves for 24 hours in the dark and at low temperature (4°C). RWC was calculated as :

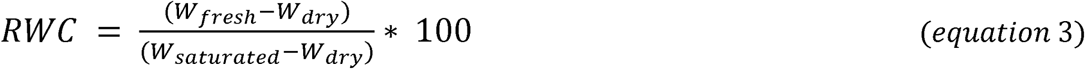

### Morpho-anatomical traits

Leaf morphological traits included total leaf area (LA_T_, cm^2^), leaf area LA (cm^2^), specific leaf area (SLA, cm^2^.g^-1^), and leaf thickness (LT, mm). Fresh leaves were scanned and analyzed with the ImageJ software (Schneider *et al*., 2012) to estimate LA and LA_T_. The scanned area and the dry masses (digital balance at a 0.0001 g precision (Mettler Toledo, Switzerland) were used to calculate SLA. A micrometer (Digit Outside Micrometre 193-101, Mitutoyo, Japan) was used to obtain LT. The root/shoot ratio of each seedling was also calculated based on dry masses.

### Microbial communities sampling

To focus on the endophytic communities inside foliar tissues, leaves were harvested at key sampling dates (Table S3), on five randomly chosen plants per treatment and three control plants per species. Leaves were surface sterilized (Compant *et al*., 2021) with successive baths of ethanol 75% (30s), ultrapure water (30s), 9% bleach (30s) and ultrapure water (30s x2) and a final bath in CTAB (30s) buffer (2% cetyl trimethylammonium bromide, 1% polyvinyl pyrrolidone, 100 mM Tris-HCl, 1.4 M NaCl, 20 mM EDTA). Leaf samples were then transferred in 2 ml UV-sterilized plastic tubes and filled with a sterile CTAB buffer. Solution of the CTAB buffer was also kept for subsequent DNA extraction as sterilization control. The samples were grinded using bead beating (Mixer mill MM 200, Retsch GmbH, Germany). For each plant individual, up to 3 replicates of 0.2 g for each sample were weighted for DNA extraction, according to the available material. Sterilization and extraction negative controls (ultrapure water) were also added. DNA was extracted using a CTAB extraction method (Carrell & Frank, 2014). We added 1400 µl of CTAB solution to 0.6 g of tissue or Whatman paper, incubated the mixture for 2 h at 60°C, and homogenized it with glass beads for 3 min. Proteins were removed by adding a two-step addition of 600 µl of chloroform, centrifuging for 10 min at 16 000 g, and isolating the top aqueous phase in a sterile tube. Nucleic acids were precipitated by adding 120 µl volume of cold 3 M sodium acetate and 1 ml cold isopropanol, froze them at -20°C for 12 h, and centrifuged for 30 min at 16 000 g. The supernatant was discarded, 700 µl of 70% ethanol was added to the solution, and centrifuged for 10 min. The air-dried pellet was resuspended with 30 µl of DNA resuspension fluid (1.0 M Tris-HCL and 0.1 M EDTA) and stored at -20°C. To characterize bacterial communities, the V5-V6 region of the bacterial 16S rDNA gene was amplified using the chloroplast-excluding forward primer 799F (Chelius and Triplett, 2001) and the reverse primer 1115R (Reysenbach and Pace, 1995). For Fungi, the ITS2 region of the rRNA gene was amplified using the ITS86F (Turenne *et al*., 1999) forward and the ITS4 (White *et al*., 1990) reverse primers as recommended by (Op De Beeck *et al*., 2014). Forward and reverse primers were tagged in 5’ with a combination of two different eight nucleotide labels. The sterilization and extraction negative controls were chosen randomly among all sterilization and extraction negative controls. Additionally, PCR negative controls (6 µL of ultra-pure water) and PCR positive controls (6 µL of 10 ng. µL^-1^ DNA, *Pseudomonas sp*. for bacteria, and *Lentinus crinitus* for fungi) were also added and amplified like all other samples. The PCR amplification was done in 50 µL and consisted of 10 µL of the Blend Master Mix 5X, 4 µL of each tagged primer (5 µM), 6 µL of DNA diluted accordingly, 24 µL of ultra-pure water and bovine serum albumin (BSA) 2 µL (10 mg.ml^-1^) to improve amplification success. PCR cycling reactions were conducted in a thermocycler (Tetrad2, BioRad, USA) with the following conditions: initial denaturation at 94 °C for 15 min, followed by 30 cycles at 94 °C for 30 sec, 58°C or 72 °C, with final extension at 72 °C for 7 min. Quality of PCR products were verified by electrophoresis using a 1 hour migration at 130 V on 1.8 % agarose gels with GelRed™ coloration (Nucleic Acid Gel Stain, 10,000X, Biotium, Glowing Products for Science™). Amplicons were then purified with magnetic beads according to the manufacturer (CleanNGS beads, CleanNA), quantified with a fluorescence-based method (Quant-it PicoGreen dsDNA assay kit ; Thermo Fisher Scientific). Amplicons replicates (1-3) of the same sample were pooled. Samples were then pooled in equimolar conditions. Two libraries were built (one for ITS2 primer and one for 16S primer) using the Fasteris MetaFast protocol (FASTERIS SA, Plan-les-Ouates, Switzerland) and sequenced on MiSeq Illumina runs using the paired-end sequencing technology (Metafast protocol, FASTERIS SA). To control for potential contaminants and false positive sequences caused by tag-switching events (Esling *et al*., 2015), the molecular experimental design comprised both sterilization, extractions and PCR negative controls that were systematically sequenced, as well as unused tag combinations.

### Bioinformatic pipelines

Sequencing reads were obtained and curated with the obitools (Boyer *et al*., 2016) and sumaclust (Mercier *et al*., 2013) for the analysis of both markers (16S and ITS2). The snakemake pipeline template is publicly hosted on github: (Anne-Sophie Benoiston. (2022). AnneSoBen/obitools_workflow: v1.0.2.GitHub.https://doi.org/10.5281/zenodo.6676577). The pipeline consisted of the following steps : 1) Merging, where forward and reverse reads were aligned to create a single consensus sequence, 2) Demultiplexing, which assigned each sequence to its sample and removed the primers, 3) Dereplication, or keeping only one representing sequence and count for strictly identical sequences, 4) Quality filtering, which removed sequences that were too short or contained ambiguous bases. The minimum sequence length accepted was 80 base pairs for ITS2 (Op De Beeck *et al*., 2014) and 100 base pairs for 16S (Redford *et al*., 2010). The similarity threshold for clustering was 97% for ITS2 and 98% for 16S. Workflows were run on the cluster of the genotoul bioinformatics platform (Bioinfo Genotoul, https://doi.org/10.15454/1.5572369328961167E12). Although the use of the OBItools allowed the removal of erroneous sequences introduced during PCR and Sequencing, further processing based on controls were used to filter spurious OTUs and contaminants using the metabaR R package (Zinger *et al*., 2020). Briefly, this additional filtering process consisted of four steps: (i) a negative control-based filtering, where MOTUs whose maximum abundance was found in extraction/PCR negative controls were removed; (ii) a reference-based filtering, where MOTUs that were too dissimilar from sequences available (< 80%) in reference databases were identified as potential chimeras and removed; (iii) an abundance-based filtering, consisting of identifying and correcting tag-jumps (Schnell *et al*., 2015) which are incorrect assignments of a few numbers of sequences corresponding to true MOTUs occurring to the wrong sample. For this, MOTUs with <0.01% of the total OTU abundance in the entire dataset were set to zero. And, finally (iv) we applied a PCR-based filter, discarding any PCR reaction that yielded fewer than 1,000 reads for the respective marker. Taxonomic assignment was done using SILVA for 16S (Quast *et al*., 2012) (Version: 1.9.10 / 1.4.9; SILVA: r138.1 -- Last Updated: 14.03.2023) and RDP classifier for ITS2 (Wang *et al*., 2007) (Version 2.11 September 2015). Finally, we checked the taxonomic assignments and kept only sequences assigned to the Bacteria and Fungi kingdoms. For the fungal dataset, 264 seedlings generated a total of 3,222,133 sequencing reads which were affiliated to fungi only and 6638 OTUs. For the bacterial dataset, 265 seedlings generated a total of 1,022,043 sequencing reads which were affiliated to bacteria only and 6149 OTUs were detected (See more details in Table S4 & S5 and rarefaction curves Figures S7 & S8). The subsequent analyses were carried out separately on the bacterial and fungal datasets.

### Analyses

#### Functional trait analyses

We used linear models to assess the influences of drought and recovery on traits separately. Treatment, species, the interaction between species and treatment and block position were set as fixed-effects terms. We preferred to fit block position as fixed effects because its number is too low to reliably estimate a variance component (Dixon, 2016; Schmid *et al*., 2017). We did not detect any significant block position effect. A Box-Cox transformation was applied to the traits prior to analysis to improve the normality of residuals.

To assess the effect of each drought and recovery treatment relative to its respective control, pairwise comparisons on untransformed data were conducted separately for each species for AGR and four selected traits, *g*_s_, *A*_sat_, *F_v_/F_m_*, and Ψ_midday_. Prior to each comparison, normality of the data was evaluated, and either Student’s *t*-tests or Wilcoxon rank-sum tests were applied accordingly.

#### Microbial analyses

The α-diversity of microbial communities, both fungi and bacteria, was estimated by computing the exponential of the Shannon index (Jost *et al*., 2010), corresponding the Hill numbers at q=1 which have been shown to provide more robust estimates of diversity (Haegeman *et al*., 2013; Calderón-Sanou *et al*., 2019). To test whether microbial diversity differed among drought treatments for each host species, we used non-parametric Kruskal–Wallis tests. When the overall test was significant (*P* < 0.05), we performed post-hoc Dunn pairwise comparisons between treatments, adjusting the *P*-values with the Benjamini–Hochberg false-discovery-rate correction. Only adjusted *P*-values were used to determine significance thresholds.

To further assess changes in community structure, we calculated two complementary β-diversity metrics at q=1: (i) dispersion, defined as the mean pairwise Hill-number distance among samples belonging to the same host species and treatment, and (ii) turnover, defined as the pairwise Hill-number distance between controls and treatment samples for each host species. We used the package HillR (Chui and Chao, 2014) and the functions hill_taxa() and hill_taxa_parti_pairwise() at q=1. For dispersion, for each host species, the mean and standard deviation of pairwise distances were calculated for each treatment and compared against the controls. For turnover, for each host species and treatment, the mean and standard deviation of pairwise distances were calculated. Control individuals were pooled across drought or recovery treatments. Treatment effects were assessed using linear mixed-effects models with random intercepts for the identity of the paired individuals. Pairwise post-hoc comparisons (among treatment for turnover, and against controls for dispersion) were obtained from estimated marginal means with Benjamini-Hochberg correction.

#### Network analyses

We performed a network analysis, using microbial metrics as traits, to investigate how the extended phenotype responded across treatments. In the network analysis, ecophysiological traits and microbial metrics are assigned as nodes and their connections as edges. We computed species mean trait values per treatment since not all traits were measured on the same individual across the experiment. We retained 14 ecophysiological traits for the analysis: *A*_sat_, *g*_s_, E, WUE, RWC, Chl, *F_v_/F_m_*, LT, LA, SLA, LA_T_, H, D, and R:S. We retained four microbial metrics for the analysis: fungal and bacterial shannon index (DivF and DivB), and fungal and bacterial dispersion (DispF and DispB). Data imputation using the *mice* package was used for two values in order to have a complete table: bacterial shannon index and bacterial dispersion index D3 for *E. falcata*.

Data was log-transformed followed by the calculation of Pearson correlation coefficients. Only correlations with p-values below 0.05 were retained; all others were set to zero. The resulting correlation matrix was used to construct a weighted, undirected graph using the *igraph* package (Csardi and Nepusz, 2005).

We extracted edge density and modularity as network-level metrics. Edge density is the ratio of observed edges to total possible edges, ranging from 0 to 1 and traits with higher edge density are considered more efficient. Modularity determines connectivity among trait modules where trait networks with higher modularity have tighter traits within than between modules. We also extracted node-level metrics such as the weighted degree, defined as the sum of all significant coefficients of correlation of a node and betweenness, which gives the number of shortest paths from all nodes to all others passing through the focal node, providing measures of a node’s importance in connecting other nodes.

To assess the significance of the observed network properties, we generated 999 null networks by randomly permuting trait values within each column of the original data matrix. For each mull network, we recalculated the correlation matrix and built the corresponding network. We computed the edge density and the modularity for each null network in order to calculate a z-score, calculated as the observed values minus the mean of values divided by the standard deviation of the null values. Two-tailed *P*-values were derived from the z-scores to assess the significance of the observed network properties relative to the null distribution.

We used R version 4.2.1 for all the following statistical analyses (R Core Team 2020).

## Results

### A. How do SF tree species recover from increasing duration of drought stress ?

#### 1. Seedling survival rates & growth

Seedlings showed very high survival throughout the experiment, with no mortality in controls, D1, or D2, while only the most severe drought (D3) resulted in notable species-specific mortality (Table S3). Survival rates after drought were lowest for *E. falcata* (6.2%), *V. surinamensis* (18.8%), *P. officinalis* (34.4%), and highest for *T. melinonii* (67.7%), *S. globulifera* (86.7%), *J. copaia* (88.5%), and *I. hostmannii* (96.8%) (Table S3). After R3 recovery, some species did not survive (*E. falcata* and *V. surinamensis*), while survival rates were low to moderate for the rest of the species: *P. officinalis* (21.9%), *T. melinonii* (25.8%), *J. copaia* (57.7%), *I. hostmannii* (58.1%) and *S. globulifera* (63.3%) (Table S3). Absolute growth rates (AGR) significantly declined with increasing drought duration, from ∼5% under D1 and D2 to 29% under D3 (Figure 2A). Growth significantly declined for 5/7 species at D3 (*t*-test, *P* < 0.02) (Figure 2A). Most species regained control-level growth after rewatering, 98.4% of control on average at R1, with only *E. falcata* differing significantly from the control (*t*-test, *P* < 0.03) (Fig 2B). Growth recovery remained incomplete under moderate drought with up to 93.6% of control on average at R2, with 3/7 species significantly differing from controls (*t*-test, *P* < 0.02) (Fig 2B). Growth recovery was lower after the most severe treatment for the remaining five species, with three species significantly differing from controls (*t*-test, *P* < 0.02), regaining only 75.5% of control at R3 (Fig 2B).

**Figure 2:**
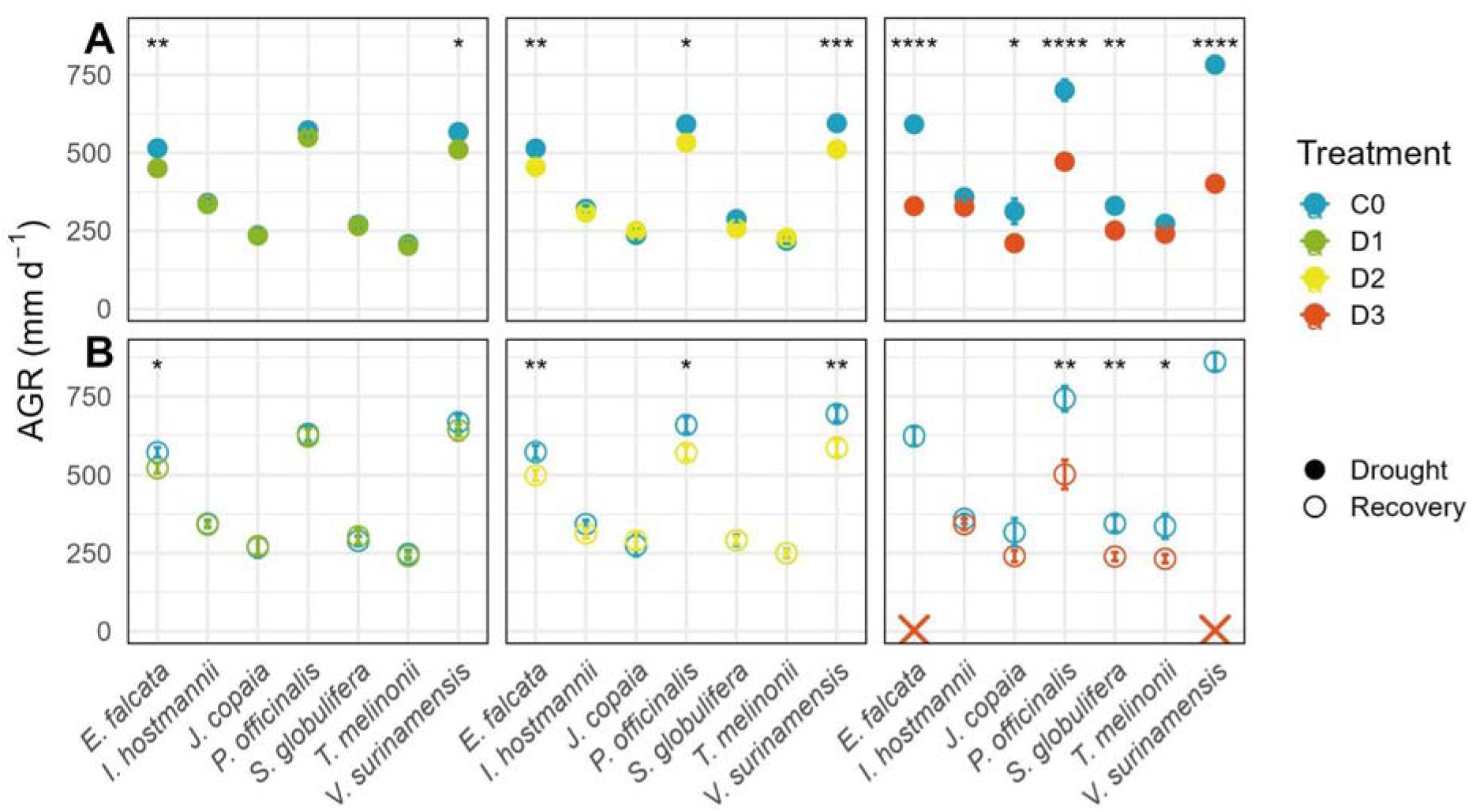
Absolute growth rate per species after drought (A) and after recovery (B). AGR is the absolute growth rate (mm d^-1^) expressed as the absolute variation in basal stem measurements between the beginning of the experiment and measurements at the end of the drought or recovery period. The red cross indicates mortality. Differences between treatment and respective control were assessed using a t.test for each species and reported as significant codes: **** P < 0.0001, *** P < 0.001, ** P < 0.01, * P < 0.05 (Table S7).

#### 2. Recovery of water- and carbon-related traits

All trait values varied significantly across the drought treatments (linear mixed effect models (LMM), *P* < 0.05, Table S8) except for LA, SLA and R:S, though the latter two exhibited significant (LMM, *P* < 0.03) species-specific drought responses (species × treatment interaction). Almost all trait values varied significantly across the recovery treatments except for *A*_sat_, WUE, RWC, LT and R:S. We did not detect any significant block position effect for any of the measured traits after drought or recovery (LMM, *P* > 0.05, Table S8). Species-specific recovery responses were found concerning *A*_sat_, *g*_s_, RWC, Ψ_midday_, Chl, *F_v_/F_m_*, LT and SLA (LMM, *P* < 0.02, Table S8). Given their strong species-specific recovery response, Figure 3 presents the variation of four key physiological traits, *g*_s_, *A*_sat_, *F_v_/F_m_*, and Ψ_midday_ values.

**Figure 3:**
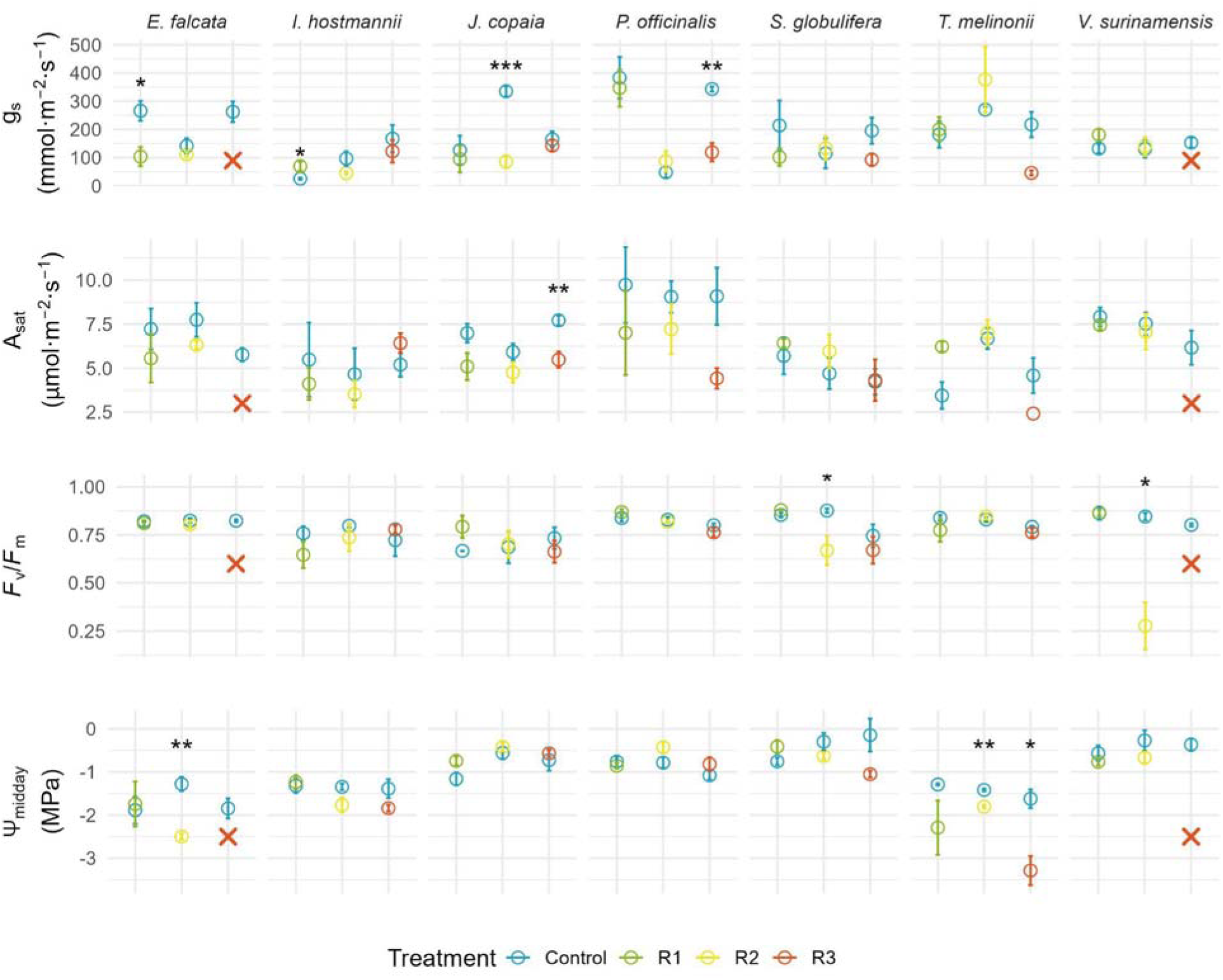
Important water and carbon related traits measured after the recovery (R1, R2, R3) with respective controls. From top to bottom: stomatal conductance (g_s_, mmol.m².s-1), carbon assimilation rate (A_sat_, µmol.m².s-1), efficiency of photosystem II (F_v_/F_m_, unitless), water potential at midday (Ψ_midday_, MPa). Statistical differences between each treatment and respective controls were assessed for each species using t-tests or Wilcoxon rank-sum tests depending on normality. The red cross indicates mortality. Significance levels are indicated by stars, **** P < 0.0001, *** P < 0.001, ** P < 0.01, * P < 0.05. Values are means ± SE.

Plant species differed in their *g*_s_ recovery responses (Figure 3). Indeed, species had closed their stomata at different time points during the initial of the drought period, with earlier stomatal closure for *E. falcata, P. officinalis, V. surinamensis* at 21 days was found, contrasting to later stomatal closure for *I. hostmannii, J. copaia, S. globulifera, T. melinonii* at 71 days (Figure S8). Upon recovery, species had similar *g*_s_ values to control except for lower *g*_s_ values than controls at R3, though only significant for *P. officinalis.* We did notice a strong variability of *g*_s_ values for the control plants that could be due to individual plant variability and day variation of climate parameters (Figure S7). Recovering *A*_sat_ values were close to control levels in *E. falcata* and *V. surinamensis*, while it even exceeded control values in *S. globulifera,* and *T. melinonii* (R1 and R2) (Figure 3). At R3, *J. copaia, P. officinalis, T. melinonii* showed incomplete recovery of photosynthetic capacity, though only significant differences were found for *J. copaia*. In contrast, for *I. hostmannii*, treatment values at R3 tended to exceed the values of the controls. Recovering *F_v_/F_m_* values were not significantly different from control values except for *S. globulifera* and *V. surinamensis* (R2). All species expressed Ψ_midday_ close to control values, except for *E.falcata and T. melinonii* at R2 and R3.

### B. How do leaf microbial communities of SF tree species recover from increasing duration of drought stress?

#### 1. Changes in microbial taxa

##### Bacterial community

We identified a total of 34 phyla and 170 orders in the leaf bacterial community of all host species across all drought treatments. The bacterial community was dominated by Proteobacteria (43-52%), Firmicutes (20-25%), Actinobacteriota (15-25%) and Bacteroidota (6-11%), with only modest shifts in their relative abundances across drought duration. Following rewatering, we identified a total of 34 phyla and 175 orders with the bacterial community dominated by Proteobacteria (46-54%), Firmicutes (25-29%), Actinobacteriota (10-15%) and Bacteroidota (6-10%). *Burkholderiales* order had the highest relative abundance in both drought and recovery treatments (31% (C0) ; 24% (D1), 32% (D2); 36% (D3), 35% (R1), 34% (R2); 28% (R3)) (Figure S9A).

##### Fungal community

We identified a total of 12 phyla and 133 orders in the leaf fungal community of all host species across all drought treatments. The community was strongly dominated by Ascomycota (80–84%) and Basidiomycota (9–14%), with 6–10% unclassified sequences. After rewatering, we identified a total of 9 phyla and 137 orders. Ascomycota remained dominant (75-84%), followed by Basidiomycota (9-18%) and 7-12% unclassified sequences. Proportions of *Xylariales* decreased with drought duration (5% (C0) ; 4% (D1), 2.3% (D2); 2% (D3)). Some orders were less abundant in recovery treatments : *Botryosphaeriales* (9-16% after D1, D2 and D3 compared to 2-5% in R1, R2 and R3) and *Trichosphaeriales* (7-10% after D1, D2 and D3 compared to 5-7% in R1, R2 and R3). Higher proportions of *Eurotiales* were recorded in all recovery treatments compared to drought (from 4-7% D1-D3 to 7-14% R1-R3) as well as *Capnodiales* in R2 (from 7% (D2) to 13%) and R3 (from 5 (D3) to 10%) (Figure S9B).

#### 2. Diversity in disturbed and recovering microbiomes

Drought did not result in a reduction of diversity in the phyllosphere (Figure S10). Bacteria diversity increased after D2 (Figure S10A), while fungal diversity remained stable across drought duration (Figure S10B). After rewatering plant host species, fungal diversity did not vary, but bacterial diversity patterns depended on the duration of drought stress period (Figure S10). Bacterial diversity significantly differed from control in R1 and R2.

Across all host species, bacterial diversity showed no significant differences among drought treatments (Kruskal–Wallis tests, all *P* > 0.05; Figure S11A). For fungal communities (Figure S11B), we only found a significant drought effect in diversity in 2/7 species (*P. officinalis* and *S. globulifera*).

Rewatering after drought induced heterogeneous responses in leaf bacterial diversity, with no single consistent pattern across host species (Figure 4A). *J. copaia* showed the strongest effects: diversity was higher at R2 compared to C0 and R1 (Dunn adj. *P* = 0.07), and significantly higher than R3 (Dunn adj. *P* = 0.03). A similar pattern was observed in *P. officinalis*, where diversity dropped at R1, peaked at R2 and decreased at R3 (although differences were not statistically significant). *S. globulifera* followed the same pattern, with lower diversity at R1, a significant increase at R2 (adj. *P* = 0.04), and lower diversity at R3. A similar trend was observed for *V. surinamensis,* with higher bacterial diversity at R2 (although not significant). Overall, intermediate drought (D2) triggered the highest bacterial diversity during recovery (R2), while the strongest drought (D3) suppressed this effect during recovery (R3).

**Figure 4:**
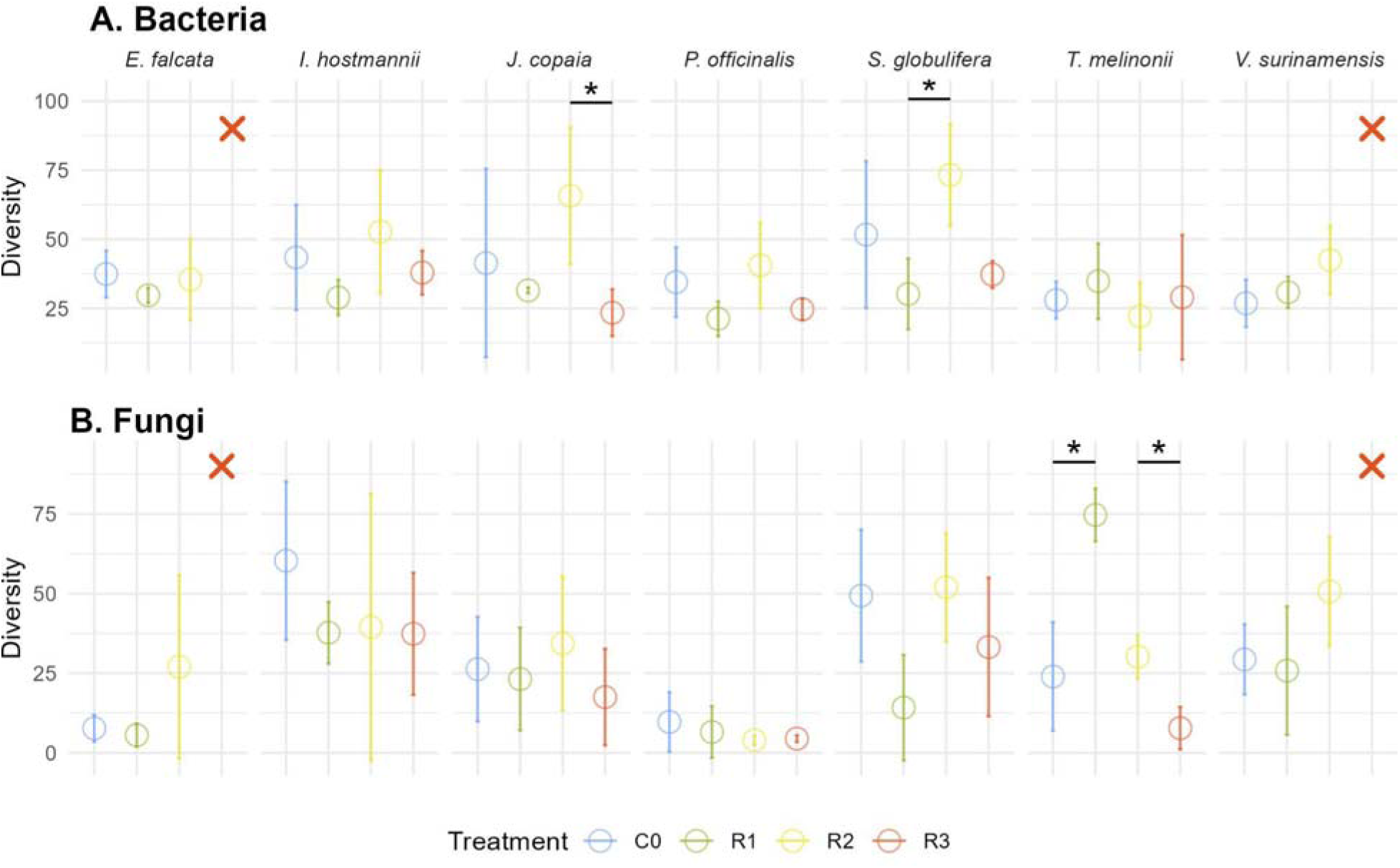
Diversity of leaf bacterial (A) and fungal (B) communities after recovery. Diversity was estimated as the exponential of Shannon entropy (Hill number at q =1). Points represent mean ± SD for each treatment. Kruskal–Wallis tests were performed to assess overall differences among treatments, followed by post-hoc Dunn pairwise comparisons with Benjamini–Hochberg correction. Asterisks indicate significant differences. The red cross indicates mortality of host plant species.

Rewatering after drought produced modest shifts in foliar fungal diversity across hosts, with significant effects detected only in *T. melinonii* (Figure 4B). In *T. melinonii*, diversity increased at R1, reaching values more than twice those of the control, before declining at R2 and dropping to very low levels at R3. *S. globulifera* and *V. surinamensis* exhibited marginal trends (Kruskal–Wallis *P* = 0.06 and *P* = 0.09, respectively), characterized by reductions at R1 and an increase at R2.

#### 3. Dispersion and turnover

Bacterial turnover during drought remained low (1.08–1.44), where 1 indicates identical community composition and higher values reflect progressively greater dissimilarity (Figure S12A). Bacterial turnover tended to increase with increasing drought duration in 5/7 host species, though not significant for most. Significant effects of drought duration were detected only in *P. officinalis* between D1-D3 and D2-D3, with higher bacterial turnover in D3 (LMM, *P* < 0.05). Marginal bacterial turnover was detected between D1-D2 in host species *E. falcata* and *T. melinonii*, with lower bacterial turnover at D2 (LMM, *P* = 0.06).

During recovery, bacterial turnover remained similarly low (1.08–1.34) and no significant differences were detected among host species, except only a marginal effect for *S. globulifera* between R1-R2 (LMM, *P* = 0.06) (Figure 5A).

**Figure 5:**
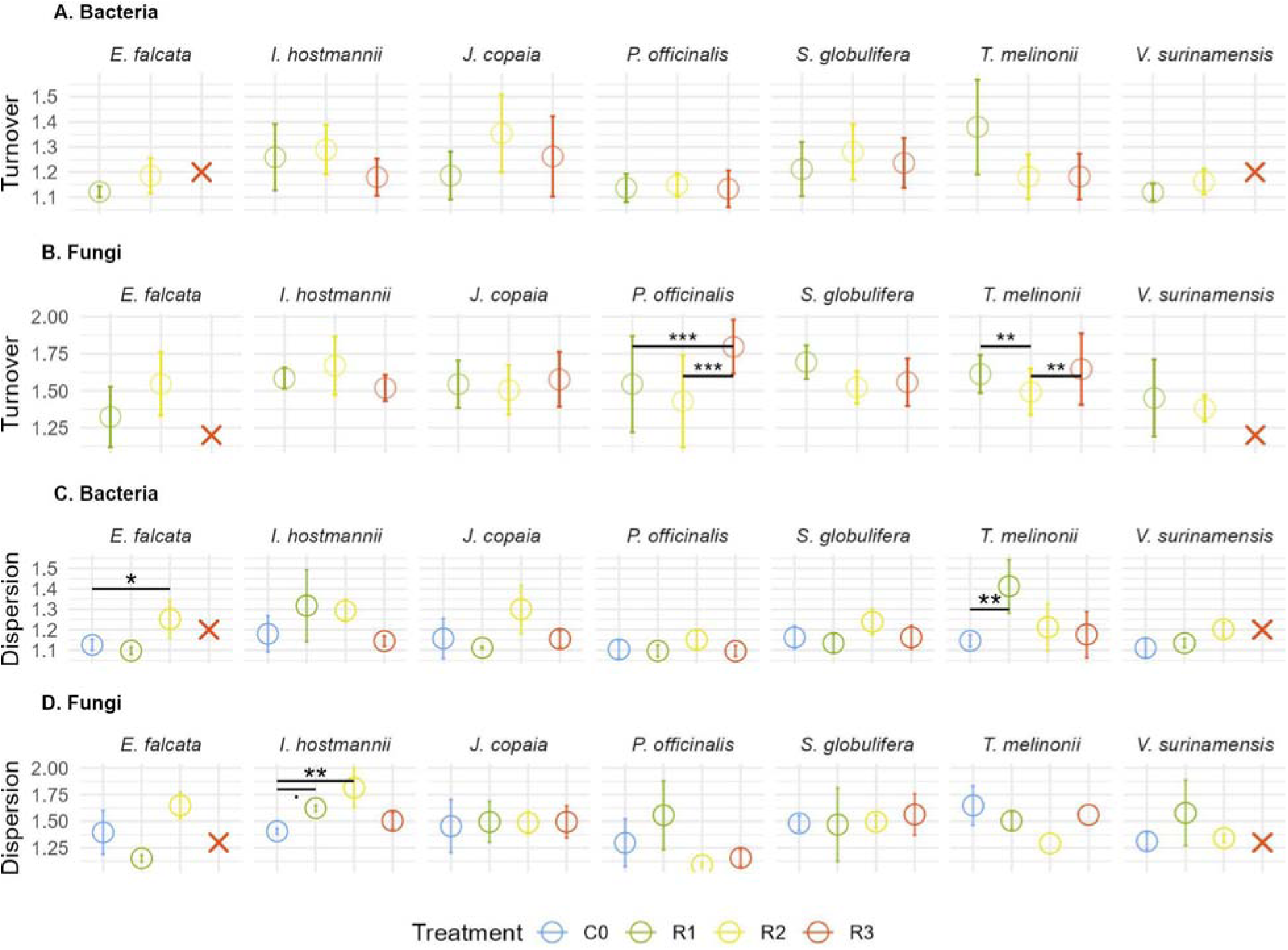
Turnover (A, B) and dispersion (C,D) of leaf bacterial and fungal communities after recovery. Turnover was calculated as pairwise β-diversity (q=1) between control and recovery treatment within the same host species using Hill distances. Dispersion or within-group β-diversity (q=1) was estimated from pairwise Hill distances among samples belonging to the same species and treatment. Control individuals were pooled across recovery treatments. Treatment effects were assessed using linear mixed-effects models with random intercepts for the identity of the paired individuals. Pairwise post-hoc comparisons (among treatment for turnover, and against C0 for dispersion) were obtained from estimated marginal means with Benjamini-Hochberg correction. Points represent mean ± SD. Asterisks indicate significant post-hoc differences. The red cross indicates mortality of host plant species.

Fungal communities showed consistently higher turnover than bacteria during drought (1.27–1.64), indicating stronger compositional divergence from controls across host species (Figure S12B). Significant drought effects were detected in 2/7 host species and. In *I. hostmannii*, fungal turnover decreased from D1 to D2 and D3 (*P* < 0.05). In contrast, *S. globulifera* showed a peak at D2, with D3 significantly lower than D2 (*P* < 0.01). We note marginal effects on fungal turnover in 2/7 host species (*J. copaia* (*P* = 0.05) and *P. officinalis* (*P* = 0.06)).

During recovery, fungal turnover remained high (1.12–1.79) across host species, indicating limited convergence back toward control communities (Figure 5B). In contrast to bacterial communities, fungi showed little decrease in turnover with prior drought intensity. Significant differences among recovery treatments were detected in 3/7 host species (*P. officinalis,* marginally for *S. globulifera,* and *T. melinonii*) (LMM, *P* < 0.05). In host *P. officinalis* fungal turnover showed a marked increase at R3. In *S. globulifera*, turnover decreased marginally (LMM, *P* = 0.06). In host *T. melinonii* fungal turnover decreased from R1 to R2 and increased from R2 to R3 (LMM, *P* < 0.01).

Dispersion values differed in their overall spread between kingdoms: bacterial communities occupied a comparatively narrow range (Drought : 1.07–1.78; Recovery: 1.06-1.64), whereas fungal communities exhibited a wider dispersion range (Drought: 1.03–1.98; Recovery: 1.05-1.96). Within these ranges, drought did not consistently reduce bacterial community dispersion (Figure S12C), instead, responses were species-specific.

Recovery trajectories also differed among host species (Figure 5C). Bacterial communities within *E. falcata*, which showed opposing shifts during drought (increased dispersion at D1, but reduced at D2), displayed a significant increase in dispersion at R2 relative to both C0 (*P* < 0.05) and R1 (*P* < 0.05), while R1 itself did not differ significantly from C0. Bacterial communities within *T. melinonii* showed a significant increase in dispersion at R1 (LMM, *P* < 0.05). Bacterial communities for which D3 affected dispersion (*I. hostmannii* (LMM, *P* = 0.05) and *P. officinalis* (LMM, *P* < 0.05), Figure S12C) had lower dispersion at R3, though not significant.

Fungal dispersion remained comparatively stable under drought, with only isolated host-specific deviations in *P. officinalis* and marginally in *S. globulifera* (Figure S12D). Fungal dispersion was significant only in host *I. hostmannii*, which trended downward during drought, significantly increased at R1 and R2 (LMM, *P* < 0.05).

### C. How does the extended phenotype of SF seedlings recover from increasing duration of drought stress?

The network analysis (Figure 6) shows that in the control, the traits are very connected together as the edge density is higher than in a random network (edge density : 0.163, *P* < 0.05), and modularity is low (0.393). Increasing drought duration significantly affects the edge density and modularity of the networks. Edge density decreases from D1 (0.150, *P* < 0.01) to D2 (0.092, *P* < 0.05) to D3 (0.078), D3 being no different from a random network, indicating that the network becomes more and more fragmented. Modularity is significantly lower in D1 (0.365, *P* < 0.05). Recovering networks all differ from the control, indicating that the extended phenotype does not return to the initial state after a drought. Recovering from D1, the R1 network is loosely connected, and modularity is higher, though not significant. Recovering from D2, the R2 network is denser than a random network (0.105, *P* < 0.05) and similarly denser to D2. R2 network is less clustered than a random network (0.328, *P* < 0.05) and less clustered than D2. Recovering from D3, the R3 network is similar to D3 and not significantly different from a random network in terms of both density and modularity.

**Figure 6:**
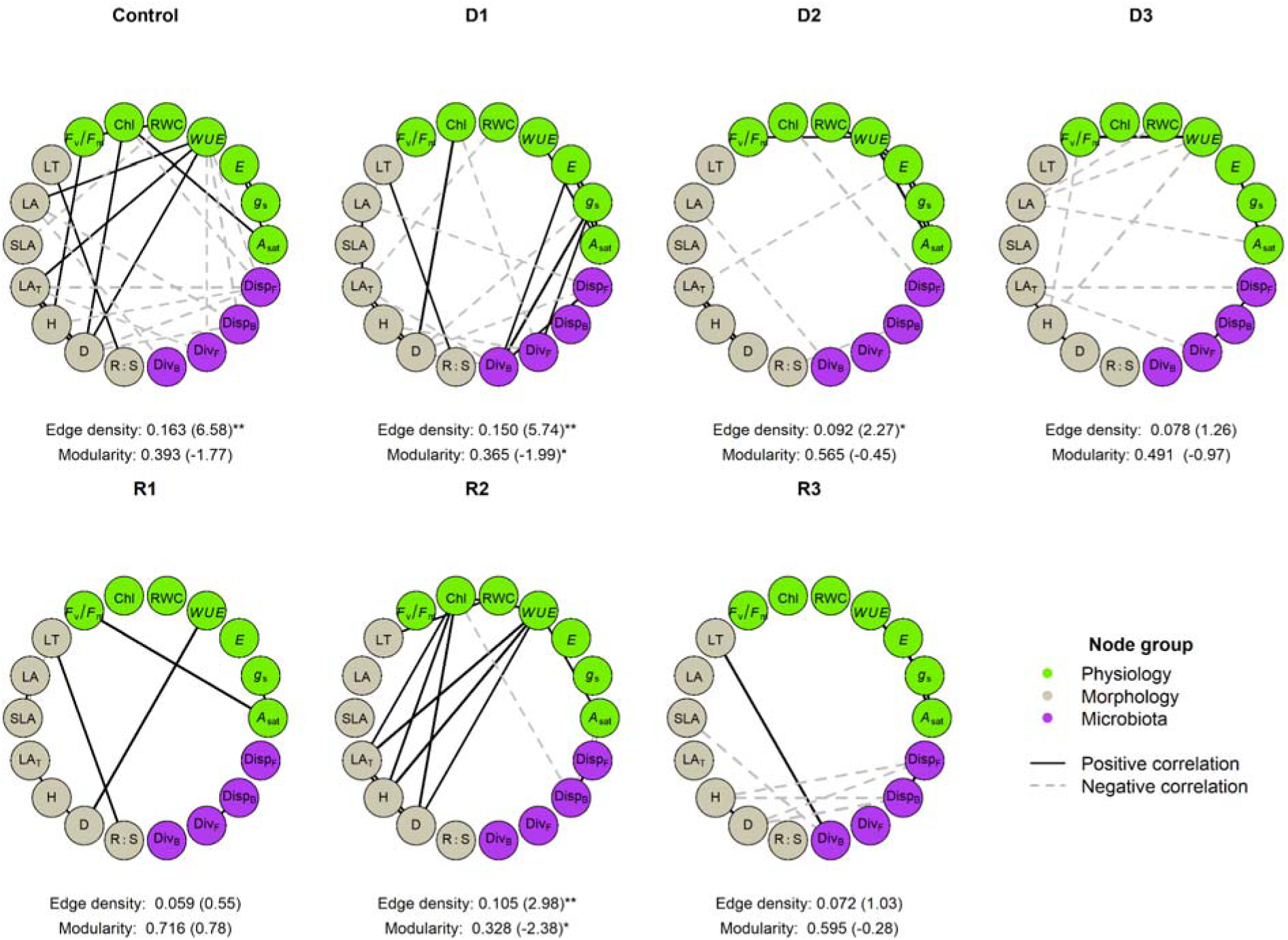
Trait correlation network shift across the control, drought (D1, D2, D3) and recovery (R1, R2, R3) treatments across all seven SF seedlings. Node color indicates functional group : green for gas exchange; grey for morphology; pink for microbiota metrics. The location of the nodes was fixed to those of the control group to allow for visual comparison. Edges represent significant Pearson correlations (P < 0.05) and correlations for which the absolute value of the correlation coefficient is set at |r| ≥ 0.2, with grey dashed indicating negative correlations and solid black edges indicating positive correlations. Edge thickness is proportional to correlation strength. Traits include LA, leaf area; SLA, specific leaf area, LA_T_, total leaf area; Chl, leaf chlorophyll content; F_v_/F_m_, the maximum quantum efficiency of photosystem II photochemistry; E, evapotranspiration, WUE, water use efficiency, A_sat_, light-saturated photosynthesis; RWC, relative water content; g_s_, stomatal conductance; LT, leaf thickness, R:S, the root to shoot ratio, H, height and D diameter, DivB and DivF, the Shannon entropy (Hill number at q =1) for bacteria and fungi respectively, DispB and DispF, beta-diversity dispersion for bacteria and fungi respectively. Each network has n=7 species except for R3 n=5 species. Network metrics are beneath each network, with observed value (z-score) significance vs. null model (999 permutations). Significant z-scores are marked with * P < 0.05 ** P < 0.01. Modularity is calculated with the Louvain algorithm.

At the node-level, two centrality measures were calculated, the weighted degree (Table S9) and betweenness (Table S10). Nodes with higher weighted degree values play a central role in the network (Table S9). In the control, the weighted degree is more or less equally partitioned between morphological, physiology and microbiota dimensions. This is similar in the D1 treatment, but with a slight re-organisation of node correlations. In D1, *g*_s_ is a central node, with microbial metrics such as DivB, DivF and DispF. In D2, E becomes a central node while microbiota metrics are not. In D3, central nodes are H, LA and WUE. We notice that H and D are always important hubs, but as drought duration increases, D becomes less central. In R1, recovering from a short drought D1, morphology traits are very central. Recovery in R2, central nodes are partitioned between the three dimensions physiology (WUE and Chl), morphology (H, D and LA_T_) and microbiota (DispB). Recovery in R3, central nodes are H and D along with DispB and DispF. For betweenness measures (Table S10), in the control, many nodes from the three dimensions (physiology, morphology and microbiota) act as mediators to other parts of the network, indicating that the information flows well between all parts of the network (Chl, H and DispF). As drought increases, gas exchange traits (*g*_s_, E, WUE) have high betweenness, as well as LA and H in D3. However, microbial metrics are not key mediators during drought, nor during recovery.

## Discussion

The aim of this study was to investigate how increasing drought duration affected the recovery of tropical tree seedlings and their associated leaf microbiota. Our results reveal that both the plant physiological recovery and microbial community recovery are strongly shaped by the plant host species identity and previous drought duration. The recovery capacity of plant physiology declined with increasing drought duration with differences among plant species which were only partially predicted by their drought tolerance strategies. Bacterial dispersion and turnover responses were strongly host species-specific, without a general directional pattern across species. Fungal communities showed more stable diversity and dispersion values, but exhibited higher turnover compared to bacterial communities during recovery, with no convergence toward control composition. Finally, none of the recovery networks returned to the architecture of the control network, regardless of prior drought duration, demonstrating that the integrated extended phenotype does not recover to initial conditions even when individual plant traits return to control values.

### Physiological recovery is partially predicted by species drought tolerance strategies

Recovery processes in plants have been less studied compared to responses to drought (Blackman *et al*., 2009; Brodribb and Cochard, 2009; Manzi *et al*., 2022), even though the capacity to recover is key to understanding drought resilience (Ruehr *et al*., 2019). Among the seven studied plant species, two contrasting drought tolerance strategies were identified (*sensus* Fletcher *et al*., 2022). Three species adopted an avoidance strategy; *E. falcata*, *P. officinalis* and *V. surinamensis* were characterized by a rapid growth during periods of high water availability (Figure 2) and closed their stomata early to minimise water loss (Figure S8). This is consistent with a conservative water-use strategy, at the expense of carbon gain (Sapes *et al*., 2019). In contrast, four species showed resistance to dehydration, maintaining *g*_s_ and Ψ_midday_ close to control values under mild to moderate drought (*I. hostmannii, J. copaia, S. globulifera* and *T. melinonii*) (Figure S8).

Avoidance species (*E. falcata*, *V. surinamensis, P. officinalis*) showed the most catastrophic outcomes. No individuals of *E. falcata* and *V. surinamensis* survived to R3, and *P. officinalis* also showed low survival rates at R3 (Table S3). Damages caused by water deficit may have been too drastic to even allow for partial recovery. *V. surinamensis* already displayed significantly low *F_v_/F_m_* values at R2 (Figure 3), which could indicate irreversible photo-damage on thylakoid membranes (Chen *et al*., 2016). According to the carbon starvation theory (McDowell *et al*., 2008), the depletion of non-structural carbohydrates in response to stomatal closure could have provoked mortality. Ziegler *et al*., (2024) showed that whole plant soluble sugars concentrations were maintained or even increased during drought in most of the tropical tree species they studied, but they declined in *E. falcata*, whose starch reserve collapsed, which could explain its mortality in our study. Interestingly, *P. officinalis* showed significantly reduced *g*_s_ and A_sat_ values relative to controls at R3, yet no difference in *F_v_/F_m_*or Ψ_midday_, suggesting that its individuals had an intact photochemical and hydraulic apparatus while carbon assimilation remained suppressed.

Among the drought-resistant species, the recovery pathway was more diversified. *I. hostmannii* showed the most robust recovery. In a previous study, *I. hostmannii* was characterized by low minimal stomatal conductance (Boisseaux *et al*., 2024) which could contribute to its desiccation tolerance during drought and its capacity to recover afterwards. For most drought-resistant species, recovery at R1 and R2, led to reversible damage, since rewatering almost completely negated the difference with the control plants (Figure 3). We observed complete recovery of Ψ_midday_ (Figure 3; except for *T. melinonii* at R2) consistent with previous studies of complete hydraulic restoration (Creek *et al*., 2018; Ruehr *et al*., 2019). This might indicate that more drought-resistant tropical species could be capable of refilling embolized xylem vessels, if previously impaired (Klein *et al*., 2018). Almost all drought-resistant species recovered A_sat_ values at R1, R2 and R3 similar to control values, though still slightly below (Figure 3), potentially reflecting persistent biochemical limitations, but *S. globulifera* and *T. melinonii* showed slightly higher A_sat_ values suggesting an overcompensation for the loss of metabolic activity during drought. Differences in AGR for *globulifera* and *T. melinonii* (Figure 2) further emphasizes that carbon could have been allocated to tissue repair instead of growth. However, *J. copaia* was the exception because of its incomplete recovery of *g*_s_ at R2 and A_sat_ at R3, indicating that drought strategy cannot fully anticipate recovery outcomes.

Overall, drought-resistant species recovered better compared with drought-avoidant species. However, the traits characterizing a drought strategy are not necessarily the same governing recovery (Yao *et al*., 2024; Zlobin, 2024).

### Recovery of leaf microbial communities : a flexible bacteriome and a stable mycobiome

Plant physiological results indicated that additional factors were necessary to fully account for the diversity of the observed recovery trajectories across SF species. We explored how leaf endophytes communities responded through changes in diversity and composition during recovery. We expected a more diverse microbiota upon recovery following a diversity-reduction during drought (Figure 1). Bacterial diversity increased in most plant host species at R2 (Figure 4). This increase in diversity reflected not only higher bacterial diversity but also greater evenness with taxa abundances becoming more equal and that fewer rare taxa dominated the recovering community. This pattern is consistent with the gradual compositional shifts observed during drought, where increasing drought duration selectively enriched stress-tolerant taxa such as *Actinobacteria* and *Burkholderiales* (Figure S9A), potentially helping the community for rapid reassembly upon rewatering (Santos-Medellin *et al*., 2017; Naylor & Coleman-Derr, 2018). A high microbial diversity has been linked to a healthy state, especially studied for the human gut microbiota (Lozupone *et al*., 2012). This re-diversification signal was, however, suppressed at R3, consistent with the idea that severe drought pushes bacterial communities beyond a compositional threshold.

Fungal diversity patterns were not significantly different and remained stable during drought and recovery (Figure 4). Previous findings have suggested that fungal communities appear to have a greater tolerance toward water limitation than bacterial communities (de Vries *et al*., 2018; Cambon *et al*., 2022; Jaeger *et al*., 2024). This is explained by fungal traits conferring tolerance to water limitation including hyphal networks, osmolyte production and thicker cell walls (de Vries *et al*., 2018; Yang *et al*., 2026). In their study, Vries *et al*. (2018) revealed that in grassland mesocosms, drought could destabilize properties in soil bacterial, but not fungal. The lack of fungal diversity change could then suggest a stable mycobiome to drought. However, this apparent stability could hide a shift in activity rather than consistent taxa diversity. As suggested by Diez-Hermano *et al*. (2022), dissimilarity metrics rather than diversity is more relevant for detecting health-related shifts in fungal communities. Yet, fungal communities exhibited consistently higher turnover values than bacteria, indicating stronger compositional divergence from control communities. This elevated fungal turnover persisted during recovery, with no convergence back to the control compositions. Microbiota may not fully recover to the same pre-existing communities, but restructure their microbial communities (Santos-Medellín *et al*., 2021). Following recovery, the relative abundance of *Botryosphaeriales* dropped (Figure S9B), a known woody plant pathogens (Slippers *et al*., 2017). We also observed a high abundance of *Xylariales* and *Capnodiales* in *P.officinalis*, while *Eurotiales* were abundant in *T. melinonii* following recovery at R3 (Figure S9B). Commonly found in tropical plants (Cambon *et al.,* 2022*), Xylariales* and *Capnodiales* are also known to have antagonistic effects against pathogens (Becker and Stadler, 2021). Besides, *Eurotiales*, which mainly includes genera like *Aspergillus* and *Penicillium*, have been reported to be plant growth promoting fungi and improve the growth of roots and shoots, the production of chlorophyll for photosynthesis in crop plants (Adedayo and Babalola, 2023). These observations suggest that fungal communities, upon rewatering, restructure toward new assemblages.

### The extended phenotype recovery strategies

The plant microbiota contributes to host phenotypic plasticity, thereby impacting the plant physiology during drought (Dastogeer *et al*., 2017). Few experiments study the extended phenotype of species to drought stress (but see Yang *et al*., 2026) and, to our knowledge, none concern the recovery of the extended phenotype of tropical tree species, making this experiment unique.

At the host plant species-level, recovery trajectories were strongly linked to specific microbiota dynamics (Figure 4 and Figure 5). Better recovery (full recovery and even enhancement) was more likely linked to a drought resistance strategy combined with a flexible bacteriome, as discussed in Yang *et al*., (2026). In their study, the *Festuca* grass showed greater responsiveness to drought, exhibiting higher trait plasticity and more pronounced shifts in its microbial communities, particularly in bacteria. In our experiment, looking at *I. hostmannii,* no mortality was observed even under extreme drought D3, and a rapid return to pre-drought physiological values upon watering, coupled with a bacterial diversification in R2. In contrast, early dysbiosis, shown as persistent bacterial dispersion increase from D1 and onwards in *V. surinamensis* was associated with physiological failure. Since the seven host species showed both drought avoidance and resistance strategies, leaf associated endophytes can participate in both drought tolerance mechanisms. Avoidant species (*E. falcata*, *P. officinalis*) showed particularly distinct fungal compositions during drought, potentially because early stomatal closure restricts fungal colonisation entry points (Arnold and Engelbrecht, 2007), leaving compositional legacies that persisted into recovery.

At the network level (Figure 6), our results showed that the extended phenotype under control conditions was highly integrated, reinforcing the holobiont concept that plants and their endophytes are a functional unit of coordination (Rosenberg & Zilber-Rosenberg, 2018; Trivedi *et al*., 2022). Increasing drought duration progressively fragmented this integration by increasing modularity and decreasing density. These findings are consistent with previous results showing that environmental stress reduces the connectivity and complexity of networks (Flores-Moreno et al., 2019; Rao *et al*., 2023; Yan *et al*., 2026). The reorganisation of central nodes across drought treatments was equally informative (Table S9 and Table S10): under D1, *g*_s_ and microbial metrics (DivB, DivF, DIspF) co-dominated network hubs, indicating early co-regulation of stomatal dynamics and microbiota; under D2 and D3, hydraulic and morphological traits displaced microbial metrics from central positions, being a network-level signature of progressive host loss of control over its microbiota (Arnault *et al*., 2023). Interestingly, none of the recovery networks returned to the control architecture, regardless of prior drought duration. This is a major result which indicates that even if individual trait values appear restored, we need to consider the plant in a multidimensional way to fully understand its resilience. The irreversibility of the extended phenotype to drought might be linked to fungal composition, as fungal communities do not converge back to control composition. The irreversibility of the extended phenotype to drought could also be linked to the length of the recovery period. Indeed, previous studies have reported a completely restored photosynthetic rate at four weeks after rewatering (Gallé *et al*., 2007; Creek *et al*., 2018). However, our methodology did not account for any possible system inertia (Van Meerbeek *et al*., 2021). Drought could have lasting effects and recovery is a dynamic process by which we could have analyzed it through repeated measurements in the weeks following drought (Ingrisch *et al*., 2023). Drought recovery assessment obtained at a single point in time might potentially provide an incomplete picture of drought recovery. Nevertheless, few studies actually allow a long recovery time (Blackman *et al*., 2019; Couchoud *et al*., 2020; Manzi *et al*., 2022). In their study, Manzi *et al*. (2022) measured 3, 7 and 14 days after rewetting and still, partial recovery was found depending on the species and the trait measured. Santos-Medellin *et al*. (2021) only allowed a week of recovery to characterize post-drought bacterial microbiome. They concluded that prolonged drought (33 days) in rice plants led to a severe microbiota restructuring that persisted even after irrigation was reestablished, but allowing more recovery time could have led to different results. Eventhough we do acknowledge that one month is still a very short time in a tree’s lifespan, it is longer than most studies and we did observe high individual trait recovery rates.

### Perspectives

In our study, we focused solely on leaf traits and leaf-associated microbiota, as plants were grown in greenhouse pots (4L) which could constrain root growth and bias results (Passioura, 2006). Future analyses should also include the root compartment for a more complete understanding of the extended phenotype. Indeed, root traits are a very useful predictor of plant responses to drought, as they are in direct contact with the soil and mediate nutrients and water uptake (Lozano *et al*., 2020). The response of a decline in soil water availability also shapes the responses of the rhizosphere and the root endosphere microorganisms (Santos-Medellin *et al*., 2017).

A central question is whether the observed drought-mediated changes in the leaf-associated communities are beneficial to the host plants, particularly in coping with drought stress and re-adjusting during the recovery. Ongoing research on endophytes have revealed a deep impact on plant survival and adaptation to abiotic constraints (Hardoim *et al*., 2015; Odokonyero *et al*., 2016; Bergna *et al*., 2018; Mengistu, 2020; Oono *et al*., 2020; Grabka *et al*., 2022). Future studies on the topic are required to determine which bacteria and fungi species are beneficial or not in order to establish a more robust causal link with drought and recovery processes. We were not able to investigate the detailed response to drought and recovery of endophyte communities for each plant species due the low sampling sizes. This was due to destructive traits (microbial metrics, RWC, SLA) which were measured on different plant individuals than non-destructive traits (such as *g*_s_, *A*_sat_, *F*_v_/*F*_m_,Ψ_midday_). This unfortunately limited the network analysis, carried out on trait means rather than individual plant values. We advise future studies to anticipate their sampling protocol to match the statistical power of their analysis.

## Conclusion

In the light to upscale these results on SF forests, we can ask ourselves : how would these habitats respond to increasing droughts? The relative resilience of tree species to the projected 30% drought increase scenario, may allow us to stay optimistic about the potential near future of these forests. The differential drought sensitivity among tree species would represent an important buffer for the community-level responses by enhancing asynchrony and affecting species abundances (Costa *et al*., 2023; Cui *et al*., 2025). However, as the climate becomes more unpredictable with high frequencies of extreme events, increased exposure of plants to drought induced die back into the future is expected (Mitchell *et al*., 2014). Additionally, because microbial communities are involved in regulating plant growth, nutrient pool and its uptake, hormone metabolism, defense responses, changes in plant associated microbial communities may alter ecosystem processes (Schimel *et al*., 2007). A better understanding of forest resilience incorporating the whole tree microbiota may improve decision-making for ecosystem management to future climates.

## Author contributions

MB: investigation, conceptualization, methodology, validation, data curation, formal analysis, visualization, project administrator, funding acquisition, writing - original draft. JYG: investigation. BB: investigation. VT: investigation, methodology. AB: investigation. JC: investigation. S-OC: investigation. SC: conceptualization, investigation, supervision, writing - original draft. HS: conceptualization, supervision, writing - original draft. CS: conceptualization, methodology, investigation, supervision, writing - original draft.

## Acknowledgements

We thank the University of Guyane, CNES and CTG for a PhD grant to M. Boisseaux. We thank the financial support from the Ceba annual project DRYER, Part of an “Investissement d’Avenir’’ grant managed by the Agence Nationale de la Recherche & the Center for the Study of Biodiversity in Amazonia (CEBA, ref. ANR-10-LABX-25-01). The internationally recognized certificate of compliance constituted from information on the permit was made available to the Access and Benefit-sharing Clearing-House with the following ABS-CH Unique Identifier (UID) ABSCH-IRCC-FR-260359-1. The reference number of the permit is the following TREL2206915S/563 issued 09/05/2022 by the French Ministry of Ecological Transition. We are grateful to Pascal Petronelli and the CIRAD inventory team for their work on tree inventories and botanical identification. Special thanks go to Luna Saunier, Léa Lebert, Clarisse Pettier, Thomas Gaquière, Sylvain Schmitt, Warren Daniel and Tristan Lafont Rapnouil for their help during forest or greenhouse seedling samplings. The climate data were provided by Météo-France to the Joint Research Unit EcoFoG for research purposes through a MétéoFrance-INRAE AgroClim convention. The authors would like to also thank the CIRAD - US Analyses TA B-49/01 34398 Montpellier Cedex 5 for the soil analyses. We are grateful to the genotoul bioinformatics platform Toulouse Occitanie for providing help and computing and storage resources (Bioinfo Genotoul, https://doi.org/10.15454/1.5572369328961167E12).

## Data availability statement

Functional trait data, microbial raw and processed data as well as the scripts for the analyses have been deposited on ZENODO (https://doi.org/10.5281/zenodo.22043055).

## Supporting Information

The following Supporting Information is available:

### Supporting Figures

Figure S1: Map of host tree species sampling collection spots.

Figure S2: Precipitation trends from 1955-2019 during the dry season.

Figure S3: Daily mean soil volumetric water content within plant pots across the different treatments.

Figure S4: Randomized block design.

Figure S5: Rarefaction curves for the bacterial dataset.

Figure S6: Rarefaction curves for the fungal dataset.

Figure S7: Control trait values throughout the experiment for all species. Error bars represent standard errors.

Figure S8: Important water and carbon related traits measured at the start of the experience (t0) and under drought treatments (D1, D2, D3) with respective controls.

Figure S9: Major (A) bacterial and (B) fungal orders detected in the leaves of the seven tropical tree seedlings across drought treatments.

Figure S10: Diversity of leaf bacterial (A) and fungal (B) communities across drought and recovery treatments.

Figure S11: Diversity of leaf bacterial (A) and fungal (B) communities across drought treatments per host species.

Figure S12: Turnover (A, B) and dispersion (C,D) of leaf bacterial and fungal communities after drought.

### Supporting Tables

Table S1: Number of different mother trees seedlings were collected from, for each species.

Table S2: Soil composition of greenhouse pots compared to seasonally flooded soils of Paracou.

Table S3: Survival rates of individuals belonging to seven SF tree species per treatment since the beginning of the experiment (t0).

Table S4: Organization of the seedling campaigns in the greenhouse.

Table S5: Mean number of reads, OTU and replicates of host seedlings for fungal analyses.

Table S6: Mean number of reads, OTU and replicates of host seedlings for bacterial analyses.

Table S7: Pairwise t.test comparisons of absolute growth rates (AGR, (mm d-1)) between the control (C0) and treatment groups (D1, D2, D3, R1, R2, R3).

Table S8: Significance tests for the effects of drought and recovery periods on traits measured on individual plants.

Table S9: Weighted degree per trait across treatments.

Table S10: Betweenness per trait across treatments.

