## Supplementary Figures for "Longer projected droughts will impair the recovery of tropical seedlings and their leaf microbiota"

### Supporting Figures

Figure S1: Map of host tree species sampling collection spots.

Figure S2: Precipitation trends from 1955-2019 during the dry season.Figure S3: Daily mean soil volumetric water content within plant pots across the different treatments.

Figure S4: Randomized block design.

Figure S5: Rarefaction curves for the bacterial dataset.

Figure S6: Rarefaction curves for the fungal dataset.

Figure S7: Control trait values throughout the experiment for all species. Error bars represent standard errors.

Figure S8: Important water and carbon related traits measured at the start of the experience (t0) and under drought treatments (D1, D2, D3) with respective controls.

Figure S9: Major (A) bacterial and (B) fungal orders detected in the leaves of the seven tropical tree seedlings across drought treatments.

Figure S10: Diversity of leaf bacterial (A) and fungal (B) communities across drought and recovery treatments.

Figure S11: Diversity of leaf bacterial (A) and fungal (B) communities across drought treatments per host species.

Figure S12: Turnover (A, B) and dispersion (C,D) of leaf bacterial and fungal communities after drought.

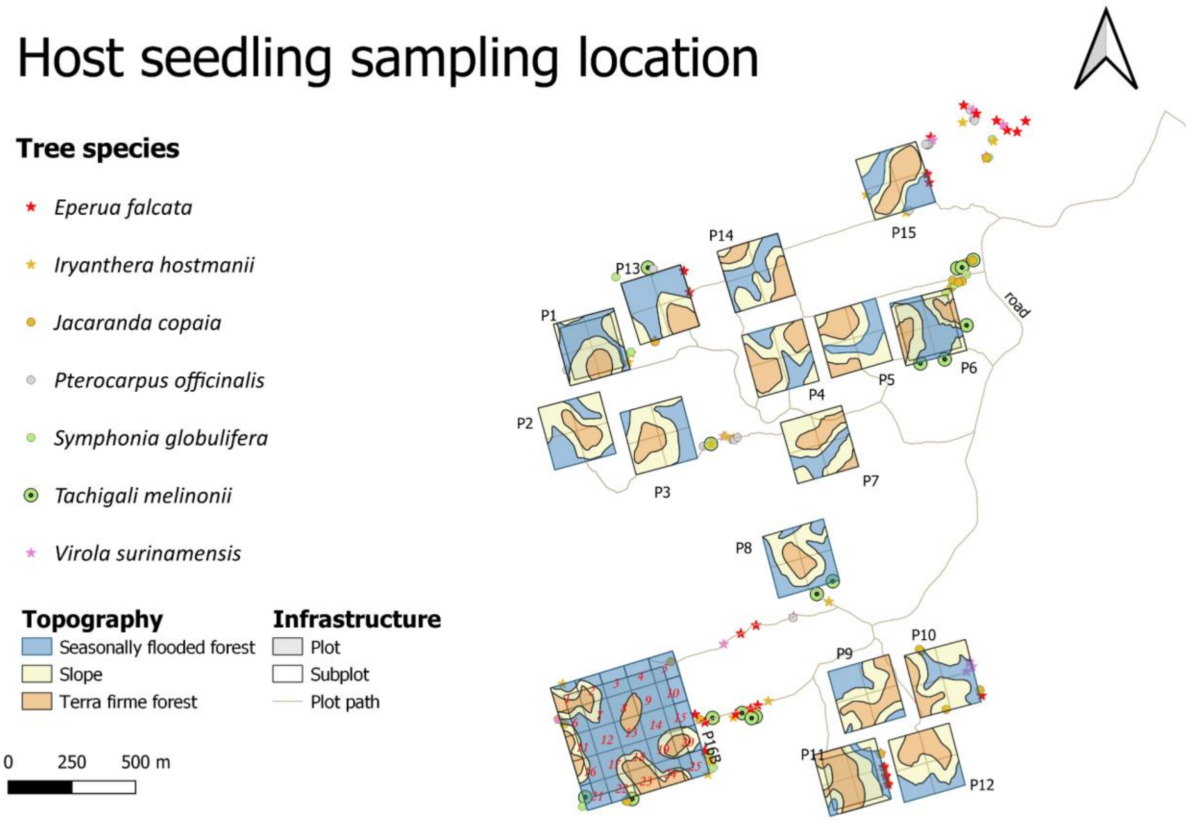


***Figure S1:*** *Map of host tree species sampling collection spots.*

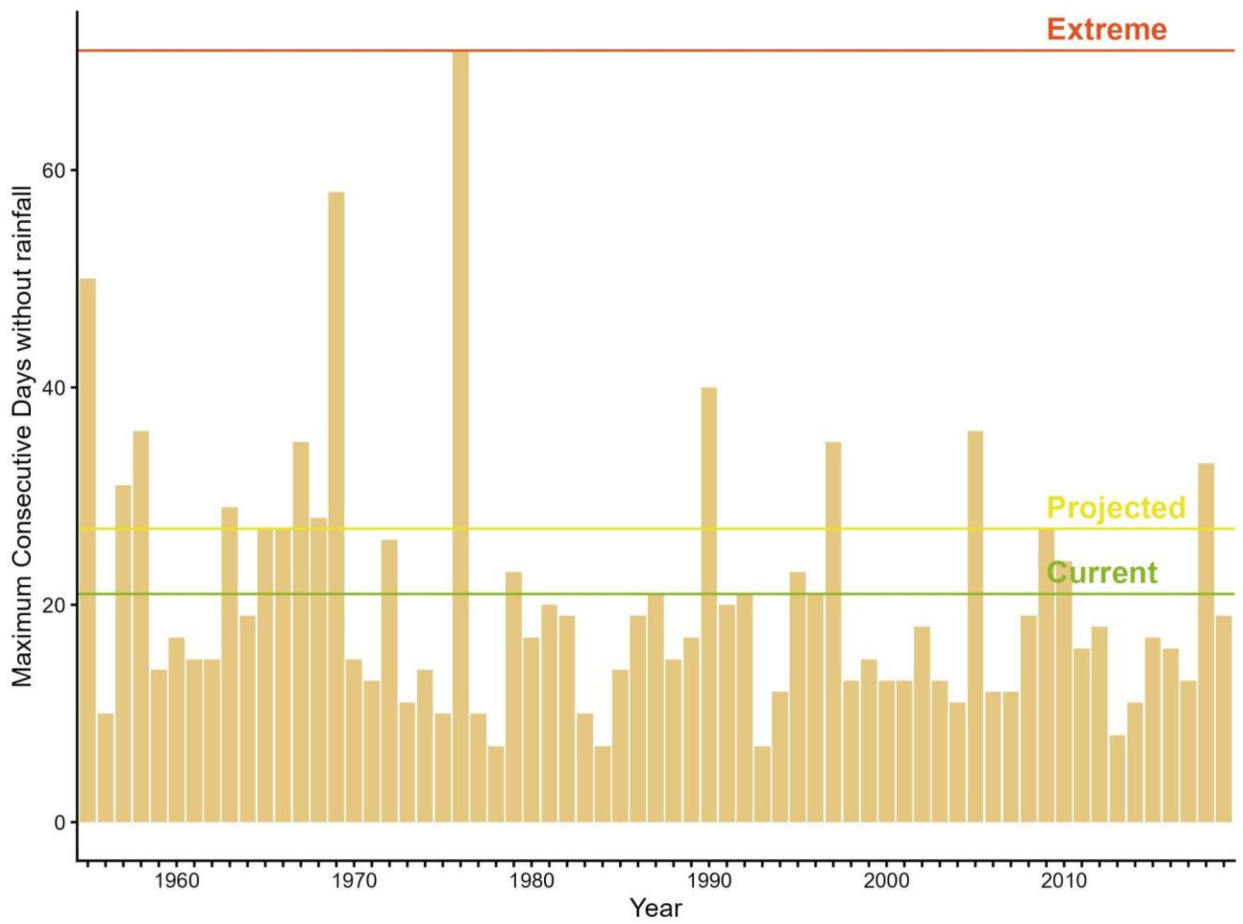


***Figure S2:*** *Precipitation trends from 1955-2019 during the dry season. The long dry season lapses from August to November (Bonal et al., 2008). Data from Meteo France Sinnamary station 97312002. Days with no rain are considered days that receive less or equal to 0.2 mm of rain. The daily rainfall records enabled us to calculate the annual mean ± SD maximum number of consecutive days without rainfall in a dry season: 20.4 ± 5.2 days. The extreme maximum number of consecutive days without rainfall was 71 days in 1976. Based on these rainfall data, we decided to simulate drought duration to match (1) the current norm of 21 days (average number of consecutive days without rainfall), (2) a projected scenario of a decrease of up to 30 % in precipitation (IPCC, 2014) 27 days (21 + 30%), and (3) an extreme event of 71 days (maximum number of consecutive dry days).*

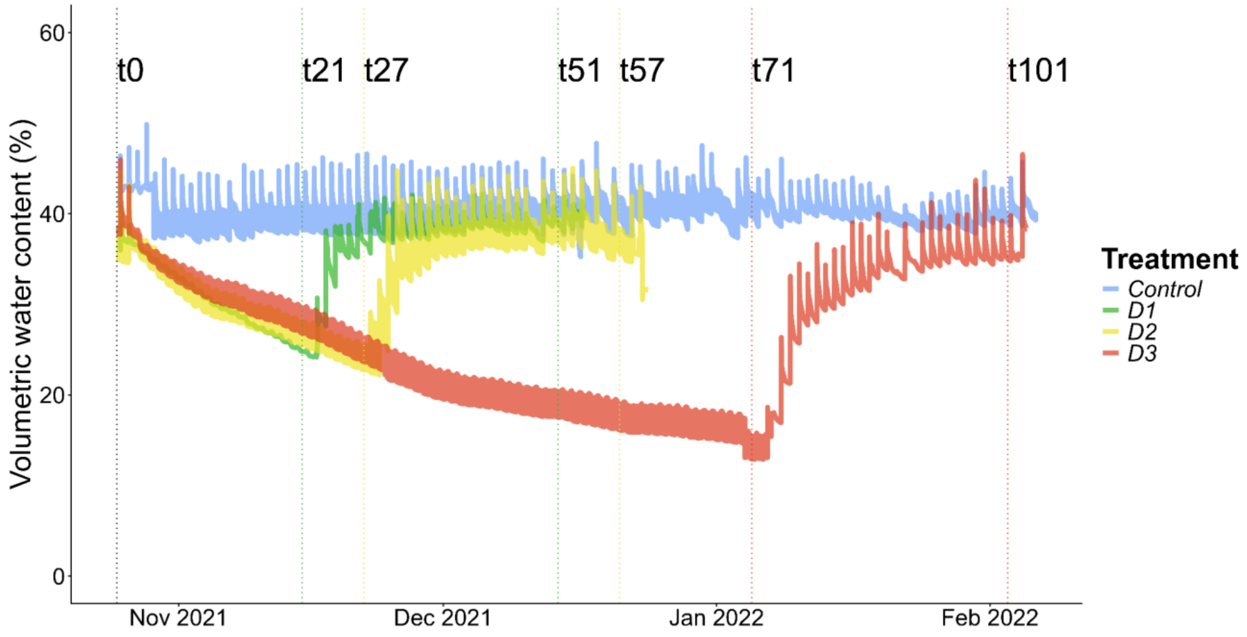


***Figure S3:*** *Daily mean soil volumetric water content within plant pots across the different treatments.*

*Soil moisture was measured using Temperature-Moisture-Sensors (Wild et al., 2019). Sensors were then calibrated with soil composition of the greenhouse pots (Table S1) to obtain the soil volumetric water content. Treatments include : control, D1, D2, D3. Sampling campaigns are shown in dotted lines, at the beginning (time t0), at the end of each drought treatment (time t21, t27, t71), at the end of each recovery period (time t51, t57, t101).*

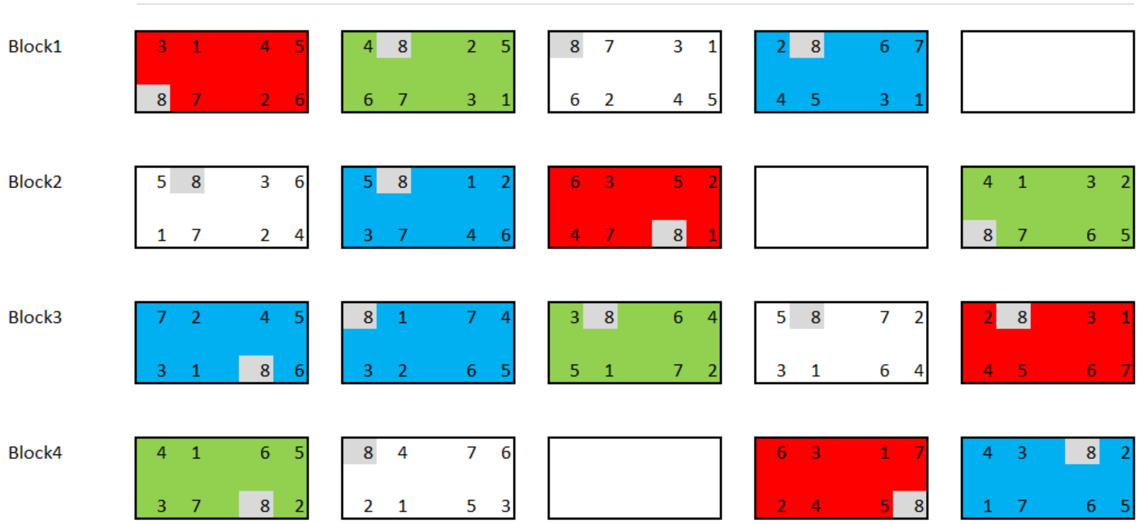


***Figure S4:*** *Randomized block design.*

*Four blocks contained the three different water treatments repeated four times, except for one control treatment repeated five times (they were more individuals for the control group). One "imaginary" (blank) treatment was repeated three times to have an even design.*

*Further notes:*

*A randomized block design is an experimental design where the experimental units are in groups called blocks. The treatments were randomly allocated to the experimental units inside each block. When all treatments appeared at least once in each block, we had a completely randomized block design. This kind of design is used to minimize the effects of systematic error. If the experimenter focuses exclusively on the differences between treatments, the effects due to variations between the different blocks should be eliminated. We used the blocksdesign R package for the construction of block and treatment designs as such:*

***blocksdesign::blocks(treatments=list(3,1,1),replicates=list(4,5,3),blocks = 4)***

*The goodness or efficiency of an experimental design can be quantified by the D-efficiency (a function of the geometric mean of the eigenvalues) and A-efficiency, a function of the arithmetic mean of the eigenvalues Both are based on the idea of average variance, as the variance gets smaller, the efficiencies go lower. The best design is the one with the highest A- and D- efficiencies. For the following block and treatment design, we obtained 0.9793704 and 0.9787234 for the A- and D- efficiencies respectively. Inside each treatment block, we organized the seven species randomly. As seven is an odd number, we used an imaginary 8th species to obtain an even design. An example is given for the control group:*

***blocksdesign::blocks(treatments=8,replicates=5,blocks = list(5,2))***

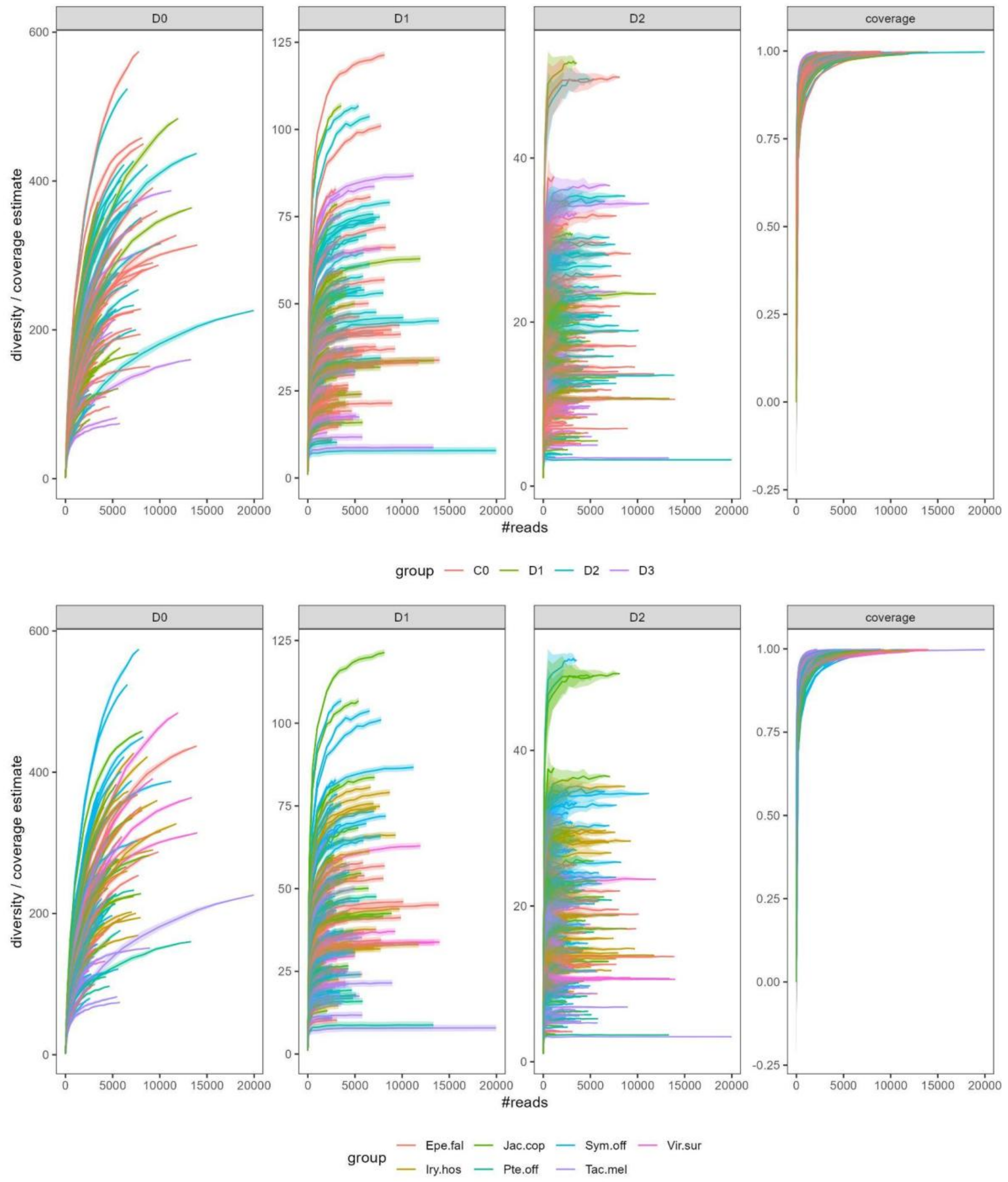


***Figure S5:*** *Rarefaction curves for the bacterial dataset.*

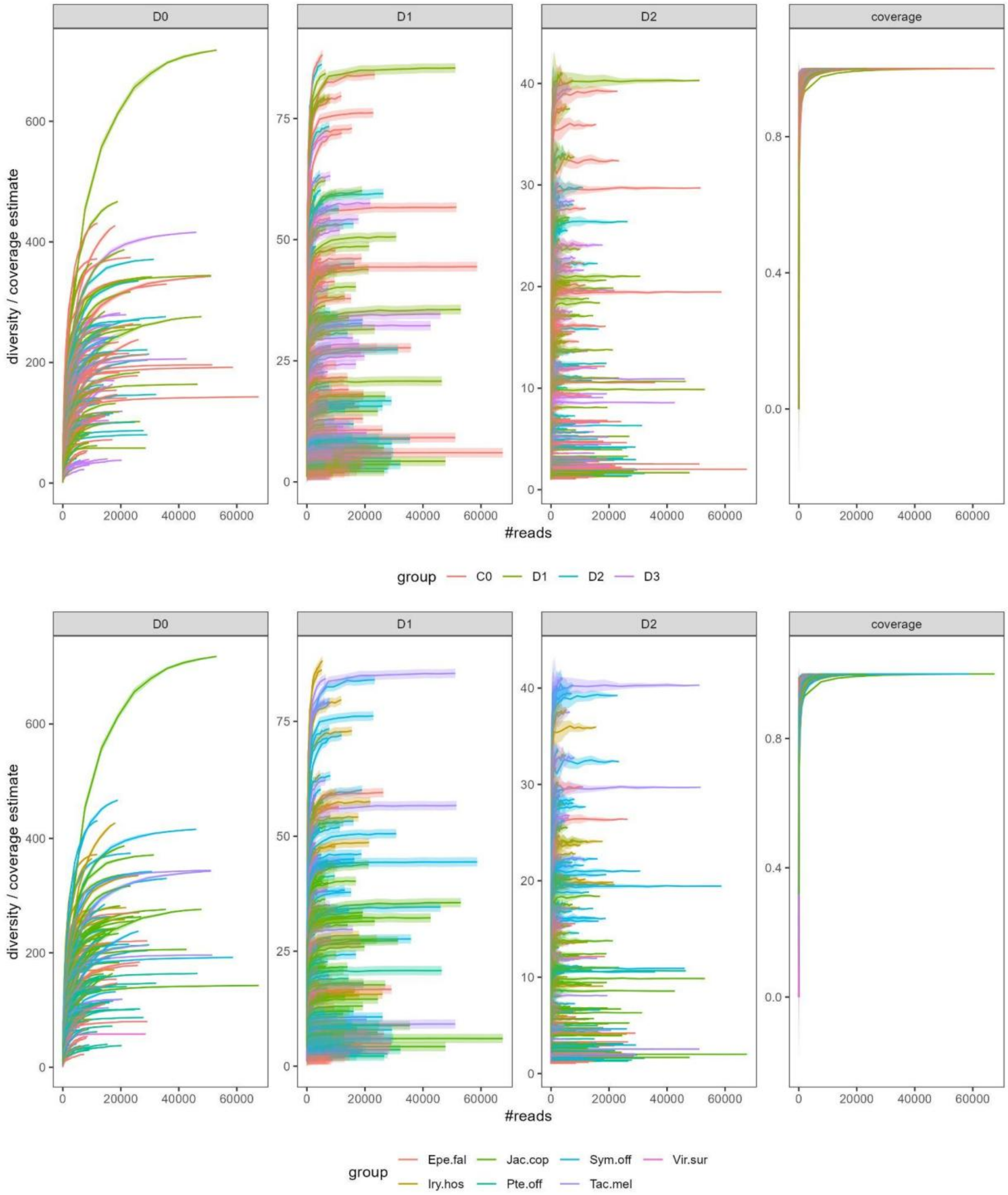


***Figure S6:*** *Rarefaction curves for the fungal dataset.*

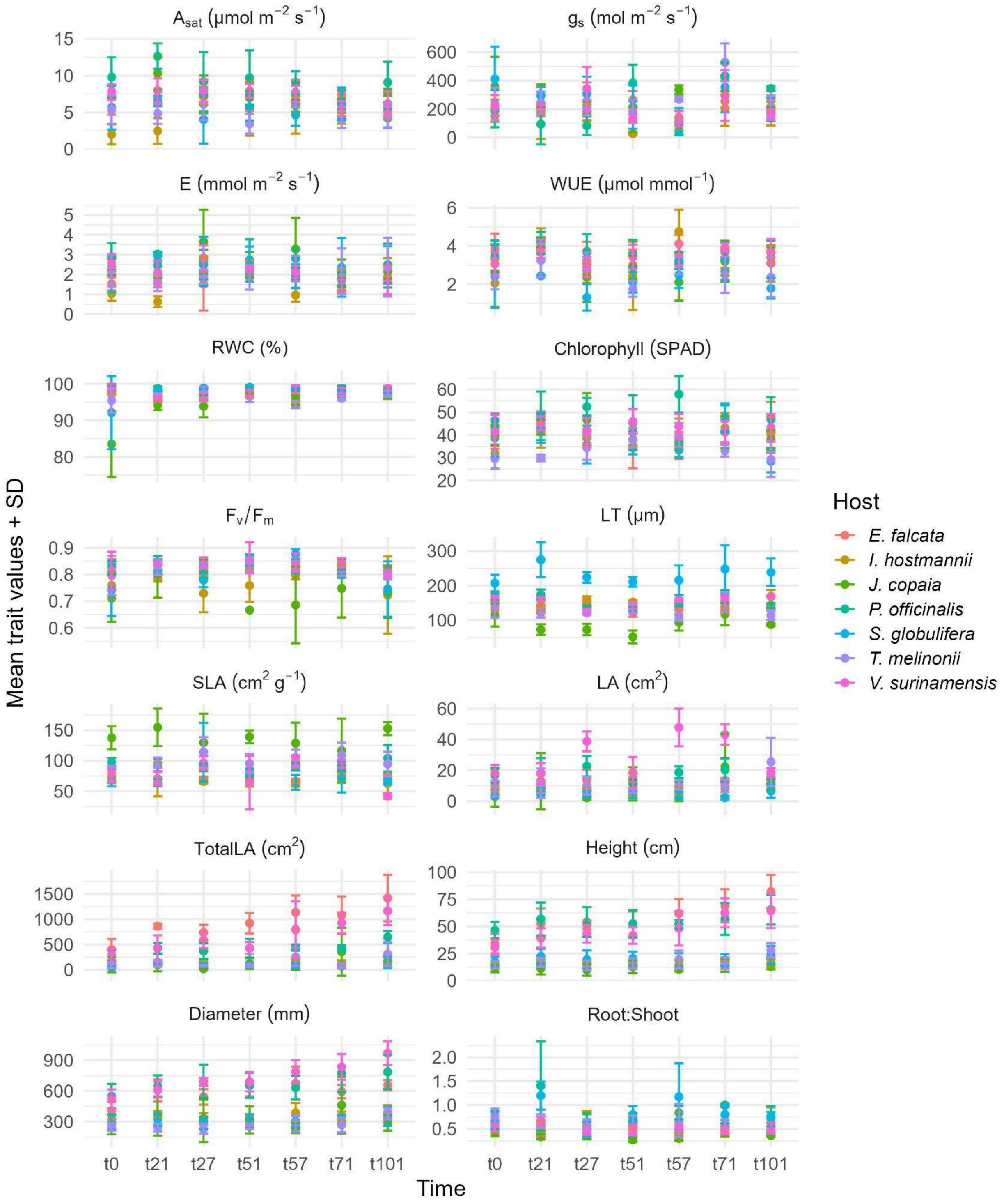


84

85 ***Figure S7:*** *Control trait values throughout the experiment for all species. Error bars represent standard*

86 *errors.*

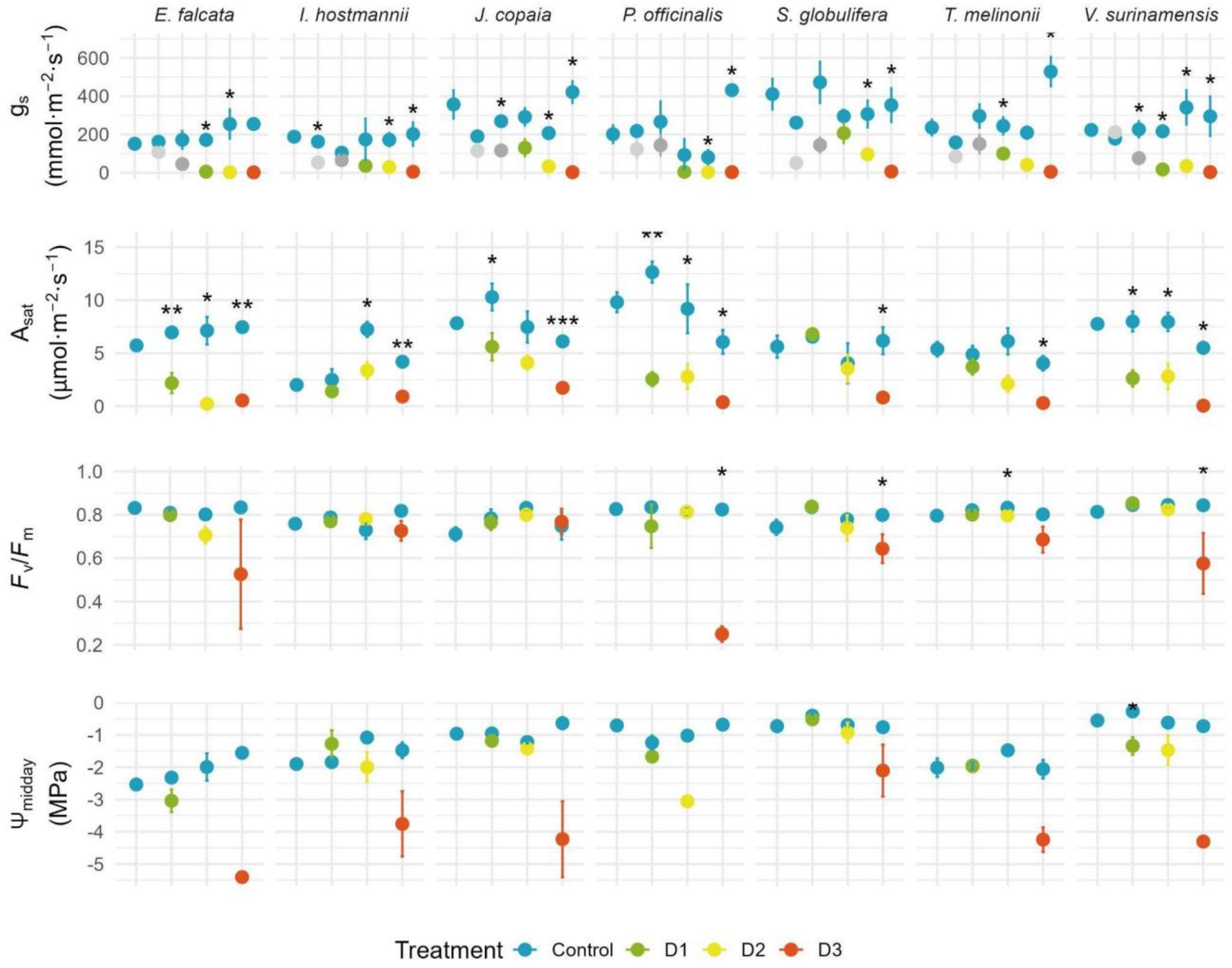


**Figure S8:** *Important water and carbon related traits measured at the start of the experience (t0) and under drought treatments (D1, D2, D3) with respective controls. From top to bottom: stomatal conductance ($g_s$, mmol.m².s$^{-1}$), carbon assimilation rate ($A_{sat}$, µmol.m².s$^{-1}$), efficiency of photosystem II ($F_v/F_m$, unitless), water potential at midday ($\Psi_{midday}$, MPa). Intermediate measurements were collected for $g_s$ at days 8 and 14. Statistical differences between each treatment and respective controls were assessed for each species using t-tests or Wilcoxon rank-sum tests depending on normality. Significance levels are indicated by stars, **** $p < 0.0001$ , *** $p < 0.001$, ** $p < 0.01$, * $p < 0.05$. Values are means ± SE.*

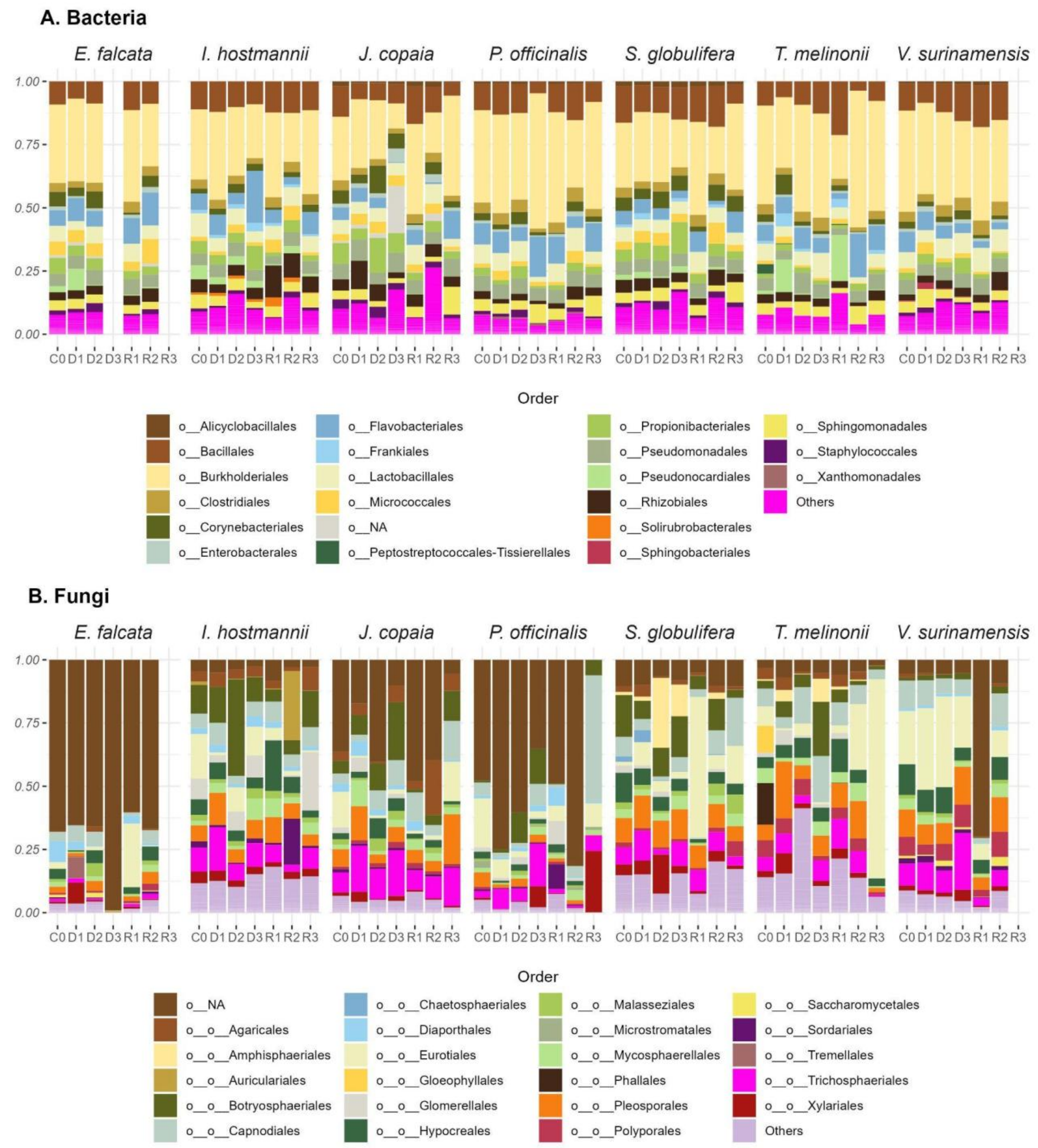


***Figure S9:*** *Major (A) bacterial and (B) fungal orders detected in the leaves of the seven tropical tree seedlings across drought treatments. NA refers to unidentified orders.*

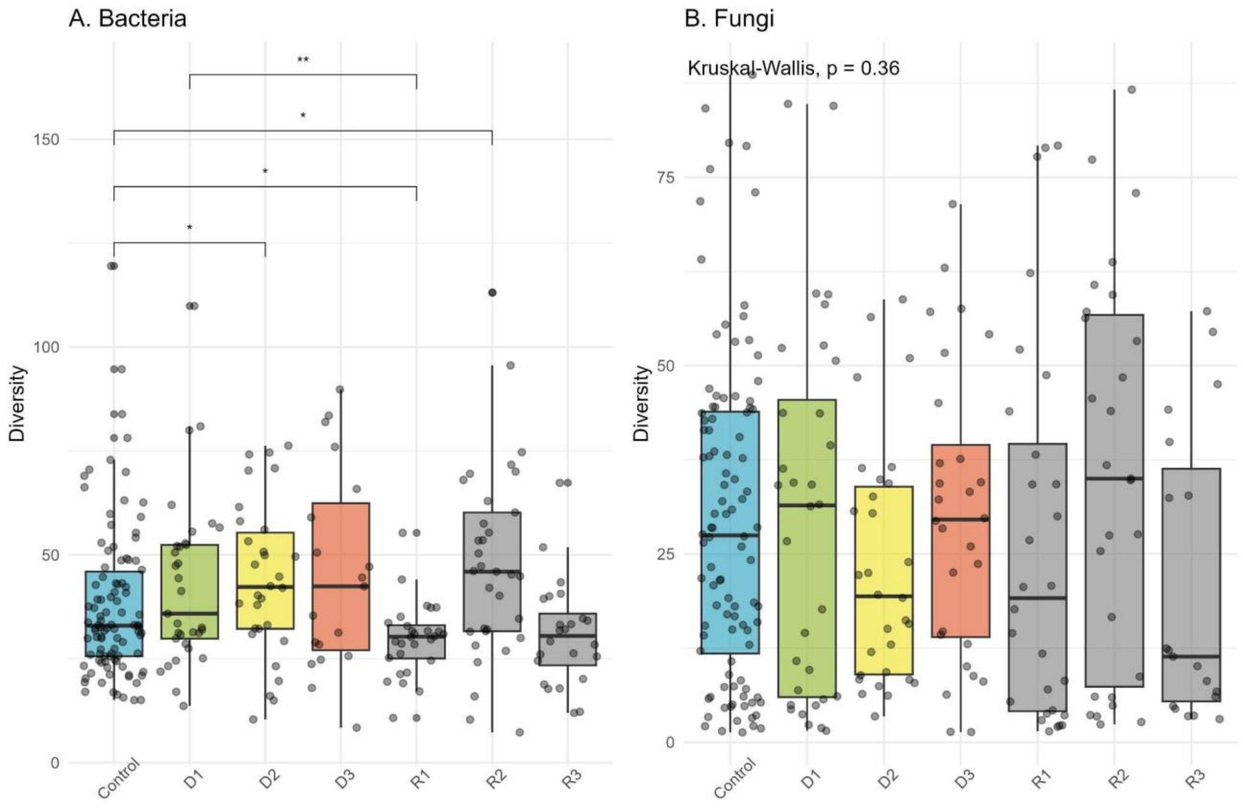


***Figure S10:*** *Diversity of leaf bacterial (A) and fungal (B) communities across drought and recovery treatments. Differences between treatments were assessed using global Kruskal–Wallis test followed by pairwise comparison using the Wilcoxon test. Only significant pairwise comparisons are shown.*

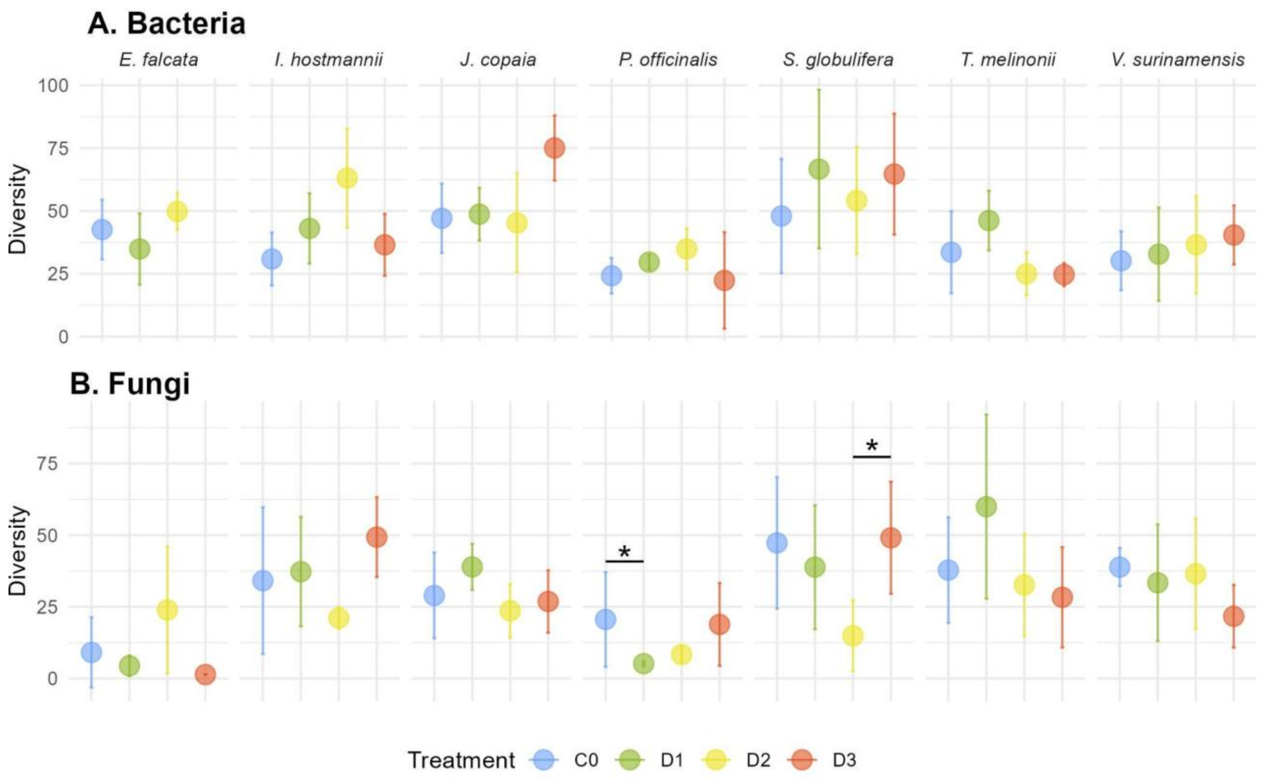


***Figure S11:*** *Diversity of leaf bacterial (A) and fungal (B) communities across drought treatments per host species. Differences between treatments were assessed using global Kruskal–Wallis test followed by pairwise comparison Dunn test. Only significant pairwise comparisons are shown. Data were insufficient to characterize bacterial diversity for E. falcata at D3.*

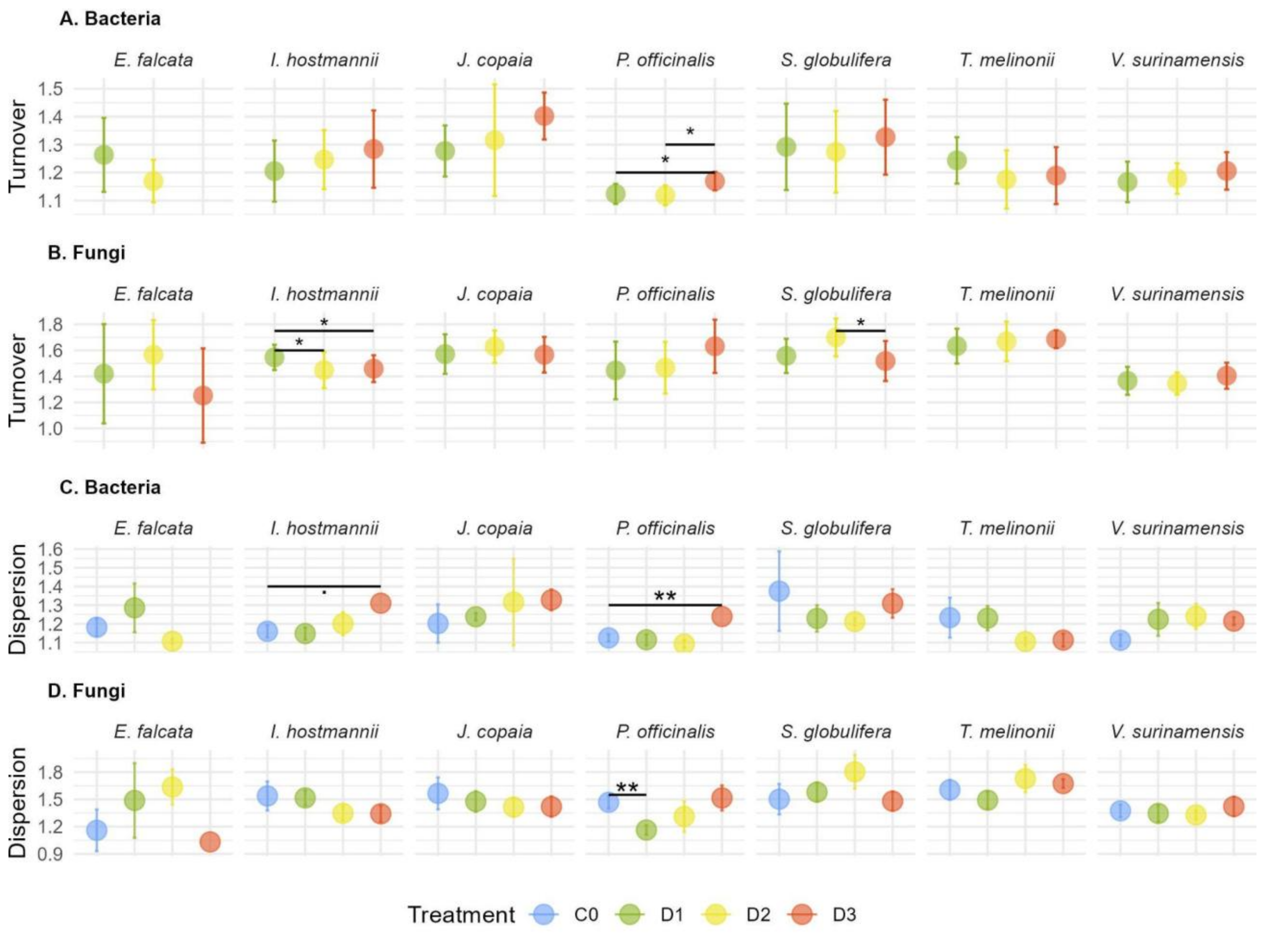


***Figure S12***: *Turnover (A, B) and dispersion (C,D) of leaf bacterial and fungal communities after drought. Turnover was calculated as pairwise β-diversity (q=1) between control and drought treatment within the same host species using Hill distances. Dispersion or within-group β-diversity (q=1) was estimated from pairwise Hill distances among samples belonging to the same species and treatment. Control individuals were pooled across drought treatments. For both metrics, treatment effects were assessed using linear mixed-effects models with random intercepts for the identity of the paired individuals. Pairwise post-hoc comparisons (among treatment for turnover, and against C0 for dispersion) were obtained from estimated marginal means with Benjamini-Hochberg correction. Points represent mean ± SD. Asterisks indicate significant post-hoc differences (P < 0.05); dots (.) indicate marginal significance (P < 0.1). Data were insufficient for bacterial microbiota of E. falcata at D3.*
