## Supplementary Tables for "Longer projected droughts will impair the recovery of tropical seedlings and their leaf microbiota"

### Supporting Tables

Table S1: Number of different mother trees seedlings were collected from, for each species.

Table S2: Soil composition of greenhouse pots compared to seasonally flooded soils of Paracou.

Table S3: Survival rates of individuals belonging to seven SF tree species per treatment since the beginning of the experiment (t0).

Table S4: Organization of the seedling campaigns in the greenhouse.

Table S5: Mean number of reads, OTU and replicates of host seedlings for fungal analyses.

Table S6: Mean number of reads, OTU and replicates of host seedlings for bacterial analyses.

Table S7: Pairwise t.test comparisons of absolute growth rates (AGR, (mm d-1)) between the control (C0) and treatment groups (D1, D2, D3, R1, R2, R3).

Table S8: Significance tests for the effects of drought and recovery periods on traits measured on individual plants.

Table S9: Weighted degree per trait across treatments.

Table S10: Betweenness per trait across treatments.

**Table S1:** Number of different mother trees seedlings were collected from, for each species.

| Species | Number of mother trees |
|---|---|
| *E. falcata* | 20 |
| *I. hostmannii* | 13 |
| *J. copaia* | 20 |
| *P. officinalis* | 9 |
| *S. globulifera* | 11 |
| *T. melinonii* | 12 |
| *V. surinamensis* | 12 |

**Table S2:** Soil composition of greenhouse pots compared to seasonally flooded soils of Paracou.

Soil composition of the greenhouse pots compared to seasonally flooded soils of Paracou.

Greenhouse and Paracou are composite samples n = 8; 8 different pots and 8 different locations in Paracou. For Paracou, soil was sampled in the 0-10 cm horizon.

| **Soil Tests** | **Greenhouse** | **Paracou** |
|---|---|---|
| Permanent wilting point (%) | 13.22 | 15.19 |
| Total carbon (%) | (0.21)<LD=0.52 | (0.22)<LD=0.52 |
| Organic matter (%) | 2.79 | 4.71 |
| Organic carbon, Corg (%) | 1.62 | 2.73 |
| Nitrogen, N (‰) | 0.69 | 1.77 |
| Corg/N | 23.54 | 15.41 |
| pH_KCl | 5.62 | 3.86 |
| pH_H2O | 6.51 | 4.67 |
| P-PO4(mg/kg)[1] | 3.72 | 5.54 |
| Ca (me/100g) | 3.53 | 0.44 |
| Mg (me/100g) | 0.33 | 0.32 |
| K (me/100g) | 0.09 | 0.07 |
| Na (me/100g) | 0.04 | 0.07 |
| Al (me/100g) | 0.01 | 0.65 |
| Mn (me/100g) | 0 | 0.01 |
| H (me/100g) | 0 | 0.27 |
| CationsEch (me/100g) | 4.01 | 1.93 |
| CEC (me/100g) | 4.2 | 2.2 |

[1] For available phosphorous, the Olsen method was used, best method for acid soils.

Results expressed in relation to the soil prepared according to the NF ISO 11464 standard, Cirad, Montpellier.

**Table S3:** Survival rates of individuals belonging to seven SF tree species per treatment since the beginning of the experiment (t0).

| Species | Control_t0 | Control_D1 | D1 | Control_D2 | D2 | Control_D3 | D3 | Control_R1 | R1 | Control_R2 | R2 | Control_R3 | R3 |
|---|---|---|---|---|---|---|---|---|---|---|---|---|---|
| *E. falcata* | 100 | 100 | 100 | 100 | 100 | 97.5 | 6.2 | 100 | 100.0 | 100 | 90.6 | 100.0 | 0.0 |
| *I. hostmannii* | 100 | 100 | 100 | 100 | 100 | 100.0 | 96.8 | 100 | 100.0 | 100 | 100.0 | 100.0 | 58.1 |
| *J. copaia* | 100 | 100 | 100 | 100 | 100 | 100.0 | 88.5 | 100 | 100.0 | 100 | 100.0 | 100.0 | 57.7 |
| *P. officinalis* | 100 | 100 | 100 | 100 | 100 | 97.5 | 34.4 | 100 | 100.0 | 100 | 100.0 | 100.0 | 21.9 |
| *S. globulifera* | 100 | 100 | 100 | 100 | 100 | 100.0 | 86.7 | 100 | 96.4 | 100 | 96.8 | 97.4 | 63.3 |
| *T. melinonii* | 100 | 100 | 100 | 100 | 100 | 100.0 | 67.7 | 100 | 100.0 | 100 | 100.0 | 100.0 | 25.8 |
| *V. surinamensis* | 100 | 100 | 100 | 100 | 100 | 100.0 | 18.8 | 100 | 100.0 | 100 | 96.9 | 100.0 | 0.0 |

**Table S4:** Organization of the seedling campaigns in the greenhouse.

| Time (days) since beginning of experiment | Details | Campaign dates |
|---|---|---|
| T0 | Beginning of experiment | 25/10/2021-28/10/2021 |
| T8 | gs measurement only | 2/11/2021-5/11/2021 |
| T14 | gs measurement only | 8/11/2021-10/11/2021 |
| T21 | Measurements on C0 and D1 | 15/11/2021-18/11/2021 |
| T27 | Measurements on C0 and D2 | 22/11/2021-25/11/2021 |
| T51 | Measurements on C0 and R1 | 14/12/2021-17/12/2021 |
| T57 | Measurements on C0 and R2 | 21/12/2021-24/12/2021 |
| T71 | Measurements on C0 and D3 | 5/01/2022-8/01/2022 |
| T101 | Measurements on C0 and R3 | 3/02/2022-06/02/2022 |

**Table S5:** Mean number of reads, OTU and replicates of host seedlings for fungal analyses.

| Species | Time | Treatment | Mean Nb reads | Mean nb Motus | n |
|---|---|---|---|---|---|
| *E. falcata* | t21 | C0 | 16676 | 250 | 2 |
| *E. falcata* | t21 | D1 | 10647 | 210 | 5 |
| *E. falcata* | t27 | C0 | 6730 | 212 | 2 |
| *E. falcata* | t27 | D2 | 12279 | 213 | 4 |
| *E. falcata* | t51 | C0 | 4650 | 113 | 3 |
| *E. falcata* | t51 | D1 | 8277 | 172 | 3 |
| *E. falcata* | t57 | C0 | 7193 | 131 | 3 |
| *E. falcata* | t57 | D2 | 8528 | 144 | 3 |
| *E. falcata* | t71 | C0 | 24276 | 350 | 3 |
| *E. falcata* | t71 | D3 | 11972 | 136 | 2 |
| *I. hostmannii* | t101 | C0 | 24841 | 347 | 3 |
| *I. hostmannii* | t101 | D3 | 12637 | 192 | 5 |
| *I. hostmannii* | t21 | C0 | 28020 | 207 | 3 |
| *I. hostmannii* | t21 | D1 | 13456 | 230 | 4 |
| *I. hostmannii* | t27 | C0 | 16623 | 216 | 2 |
| *I. hostmannii* | t27 | D2 | 8545 | 140 | 4 |
| *I. hostmannii* | t51 | D1 | 5488 | 135 | 3 |
| *I. hostmannii* | t57 | C0 | 9556 | 86 | 2 |
| *I. hostmannii* | t57 | D2 | 14371 | 164 | 4 |
| *I. hostmannii* | t71 | D3 | 18102 | 189 | 4 |
| *J. copaia* | t101 | C0 | 16201 | 177 | 3 |
| *J. copaia* | t101 | D3 | 8460 | 175 | 4 |
| *J. copaia* | t21 | C0 | 32274 | 171 | 3 |
| *J. copaia* | t21 | D1 | 11599 | 110 | 5 |
| *J. copaia* | t27 | C0 | 9605 | 187 | 3 |
| *J. copaia* | t27 | D2 | 9106 | 158 | 4 |
| *J. copaia* | t51 | C0 | 29978 | 220 | 2 |
| *J. copaia* | t51 | D1 | 12681 | 153 | 5 |
| *J. copaia* | t57 | C0 | 9747 | 114 | 3 |
| *J. copaia* | t57 | D2 | 14203 | 142 | 4 |
| *J. copaia* | t71 | C0 | 15538 | 239 | 2 |
| *J. copaia* | t71 | D3 | 13477 | 165 | 5 |
| *P. officinalis* | t101 | C0 | 6092 | 136 | 3 |
| *P. officinalis* | t101 | D3 | 14545 | 212 | 5 |
| *P. officinalis* | t21 | C0 | 2968 | 156 | 2 |
| *P. officinalis* | t21 | D1 | 10466 | 124 | 5 |
| *P. officinalis* | t27 | C0 | 6992 | 183 | 3 |
| *P. officinalis* | t27 | D2 | 6054 | 120 | 4 |
| *P. officinalis* | t51 | C0 | 6019 | 170 | 3 |
| *P. officinalis* | t51 | D1 | 17124 | 152 | 5 |
| *P. officinalis* | t57 | C0 | 4140 | 83 | 3 |
| *P. officinalis* | t57 | D2 | 7674 | 59 | 4 |
| *P. officinalis* | t71 | C0 | 3656 | 148 | 2 |
| *P. officinalis* | t71 | D3 | 10910 | 146 | 4 |

| Species | Time | Treatment | Mean Nb reads | Mean nb Motus | n |
|---|---|---|---|---|---|
| *S. globulifera* | t101 | C0 | 6726 | 67 | 3 |
| *S. globulifera* | t101 | D3 | 24234 | 278 | 3 |
| *S. globulifera* | t21 | C0 | 5410 | 115 | 3 |
| *S. globulifera* | t21 | D1 | 18882 | 213 | 4 |
| *S. globulifera* | t27 | C0 | 7704 | 109 | 3 |
| *S. globulifera* | t27 | D2 | 1930 | 103 | 5 |
| *S. globulifera* | t51 | C0 | 2337 | 122 | 1 |
| *S. globulifera* | t51 | D1 | 22936 | 203 | 4 |
| *S. globulifera* | t57 | C0 | 12298 | 105 | 3 |
| *S. globulifera* | t57 | D2 | 9680 | 183 | 5 |
| *S. globulifera* | t71 | C0 | 10817 | 117 | 3 |
| *S. globulifera* | t71 | D3 | 15907 | 196 | 5 |
| *T. melinonii* | t101 | C0 | 20378 | 160 | 3 |
| *T. melinonii* | t101 | D3 | 11534 | 100 | 2 |
| *T. melinonii* | t21 | C0 | 12122 | 162 | 2 |
| *T. melinonii* | t21 | D1 | 18756 | 116 | 4 |
| *T. melinonii* | t27 | C0 | 7173 | 136 | 3 |
| *T. melinonii* | t27 | D2 | 11767 | 210 | 4 |
| *T. melinonii* | t51 | C0 | 16309 | 174 | 3 |
| *T. melinonii* | t51 | D1 | 16233 | 184 | 4 |
| *T. melinonii* | t57 | C0 | 12216 | 165 | 3 |
| *T. melinonii* | t57 | D2 | 8374 | 287 | 2 |
| *T. melinonii* | t71 | C0 | 11319 | 151 | 1 |
| *T. melinonii* | t71 | D3 | 14553 | 145 | 3 |
| *V. surinamensis* | t21 | C0 | 12461 | 199 | 3 |
| *V. surinamensis* | t21 | D1 | 4814 | 119 | 5 |
| *V. surinamensis* | t27 | C0 | 4640 | 138 | 3 |
| *V. surinamensis* | t27 | D2 | 21100 | 191 | 5 |
| *V. surinamensis* | t51 | C0 | 7517 | 178 | 3 |
| *V. surinamensis* | t51 | D1 | 10414 | 187 | 4 |
| *V. surinamensis* | t57 | C0 | 21936 | 128 | 3 |
| *V. surinamensis* | t57 | D2 | 3030 | 117 | 5 |
| *V. surinamensis* | t71 | C0 | 6992 | 163 | 2 |
| *V. surinamensis* | t71 | D3 | 11222 | 186 | 5 |

**Table S6:** Mean number of reads, OTU and replicates of host seedlings for bacterial analyses.

| Species | Time | Treatment | Mean Nb reads | Mean nb Motus | n |
|---|---|---|---|---|---|
| *E. falcata* | t21 | C0 | 2827 | 266 | 3 |
| *E. falcata* | t21 | D1 | 4264 | 260 | 5 |
| *E. falcata* | t27 | C0 | 3392 | 192 | 3 |
| *E. falcata* | t27 | D2 | 2136 | 163 | 5 |
| *E. falcata* | t51 | C0 | 2758 | 183 | 3 |
| *E. falcata* | t51 | D1 | 5532 | 189 | 5 |
| *E. falcata* | t57 | C0 | 3055 | 197 | 3 |
| *E. falcata* | t57 | D2 | 2212 | 137 | 5 |
| *E. falcata* | t71 | C0 | 3285 | 205 | 3 |
| *I. hostmannii* | t101 | C0 | 5677 | 194 | 3 |
| *I. hostmannii* | t101 | D3 | 3968 | 255 | 5 |
| *I. hostmannii* | t21 | C0 | 2928 | 116 | 2 |
| *I. hostmannii* | t21 | D1 | 8027 | 299 | 5 |
| *I. hostmannii* | t27 | C0 | 5346 | 259 | 3 |
| *I. hostmannii* | t27 | D2 | 2372 | 183 | 5 |
| *I. hostmannii* | t51 | C0 | 3150 | 160 | 2 |
| *I. hostmannii* | t51 | D1 | 4048 | 318 | 3 |
| *I. hostmannii* | t57 | C0 | 2312 | 167 | 3 |
| *I. hostmannii* | t57 | D2 | 5732 | 192 | 5 |
| *I. hostmannii* | t71 | C0 | 1963 | 239 | 1 |
| *I. hostmannii* | t71 | D3 | 7182 | 350 | 2 |
| *J. copaia* | t101 | C0 | 3661 | 270 | 2 |
| *J. copaia* | t101 | D3 | 5206 | 283 | 4 |
| *J. copaia* | t21 | C0 | 1570 | 142 | 2 |
| *J. copaia* | t21 | D1 | 2394 | 116 | 4 |
| *J. copaia* | t27 | C0 | 7538 | 333 | 2 |
| *J. copaia* | t27 | D2 | 1923 | 170 | 5 |
| *J. copaia* | t51 | C0 | 2429 | 189 | 3 |
| *J. copaia* | t51 | D1 | 2489 | 215 | 3 |
| *J. copaia* | t57 | C0 | 5898 | 182 | 3 |
| *J. copaia* | t57 | D2 | 2627 | 190 | 5 |
| *J. copaia* | t71 | C0 | 2063 | 164 | 2 |
| *J. copaia* | t71 | D3 | 6181 | 275 | 3 |
| *P. officinalis* | t101 | C0 | 3250 | 184 | 3 |
| *P. officinalis* | t101 | D3 | 5925 | 256 | 5 |
| *P. officinalis* | t21 | C0 | 5406 | 254 | 2 |
| *P. officinalis* | t21 | D1 | 4731 | 210 | 5 |
| *P. officinalis* | t27 | C0 | 2237 | 164 | 3 |
| *P. officinalis* | t27 | D2 | 2245 | 126 | 4 |
| *P. officinalis* | t51 | D1 | 3274 | 212 | 4 |
| *P. officinalis* | t57 | C0 | 4037 | 289 | 3 |
| *P. officinalis* | t57 | D2 | 4119 | 283 | 5 |
| *P. officinalis* | t71 | D3 | 2116 | 190 | 2 |
| *S. globulifera* | t101 | C0 | 3988 | 220 | 3 |
| *S. globulifera* | t101 | D3 | 5138 | 239 | 5 |
| *S. globulifera* | t21 | C0 | 6796 | 252 | 3 |
| *S. globulifera* | t21 | D1 | 4577 | 248 | 5 |
| *S. globulifera* | t27 | C0 | 3074 | 140 | 3 |
| *S. globulifera* | t27 | D2 | 1901 | 145 | 3 |
| *S. globulifera* | t51 | C0 | 2758 | 116 | 3 |
| *S. globulifera* | t51 | D1 | 6707 | 317 | 5 |
| *S. globulifera* | t57 | C0 | 1969 | 150 | 3 |
| *S. globulifera* | t57 | D2 | 3649 | 220 | 5 |
| *S. globulifera* | t71 | C0 | 2490 | 135 | 2 |
| *S. globulifera* | t71 | D3 | 2524 | 250 | 4 |
| *T. melinonii* | t101 | C0 | 3084 | 159 | 2 |
| *T. melinonii* | t101 | D3 | 6024 | 227 | 5 |
| *T. melinonii* | t21 | C0 | 1926 | 128 | 3 |
| *T. melinonii* | t21 | D1 | 3822 | 214 | 4 |
| *T. melinonii* | t27 | C0 | 2670 | 125 | 3 |
| *T. melinonii* | t27 | D2 | 4392 | 257 | 4 |
| *T. melinonii* | t51 | C0 | 2090 | 196 | 2 |
| *T. melinonii* | t51 | D1 | 1933 | 178 | 5 |
| *T. melinonii* | t57 | C0 | 3022 | 175 | 3 |
| *T. melinonii* | t57 | D2 | 4858 | 256 | 4 |
| *T. melinonii* | t71 | C0 | 2265 | 192 | 3 |
| *T. melinonii* | t71 | D3 | 6207 | 283 | 4 |
| *V. surinamensis* | t21 | C0 | 2994 | 156 | 2 |
| *V. surinamensis* | t21 | D1 | 3292 | 164 | 5 |
| *V. surinamensis* | t27 | C0 | 3949 | 183 | 2 |
| *V. surinamensis* | t27 | D2 | 3968 | 238 | 4 |
| *V. surinamensis* | t51 | C0 | 3388 | 240 | 2 |
| *V. surinamensis* | t51 | D1 | 5048 | 226 | 3 |
| *V. surinamensis* | t57 | C0 | 2688 | 146 | 2 |
| *V. surinamensis* | t57 | D2 | 3108 | 253 | 4 |
| *V. surinamensis* | t71 | C0 | 2143 | 144 | 3 |
| *V. surinamensis* | t71 | D3 | 6175 | 258 | 4 |

**Table S7:** Pairwise t.test comparisons of absolute growth rates (AGR, (mm d-1)) between the control (C0) and treatment groups (D1, D2, D3, R1, R2, R3). P-values were calculated using two-sample *t*-tests and adjusted using the *Holm* method. Bold p-values indicate significant differences ($p < 0.05$); a p-value of 0 indicates $p < 0.001$; n is the sample size and CI 95 is the confidence interval of the difference in means.

| Species | group1 | group2 | p.value | n | CI_95 |
|---|---|---|---|---|---|
| *E. falcata* | C0 | D1 | **0.002** | 34 / 32 | [23.79, 104.18] |
| *E. falcata* | C0 | D2 | **0.005** | 32 / 32 | [19.08, 100.22] |
| *E. falcata* | C0 | D3 | **0** | 22 / 2 | [222.81, 302.76] |
| *E. falcata* | C0 | R1 | **0.03** | 28 / 27 | [5.13, 96.14] |
| *E. falcata* | C0 | R2 | **0.007** | 26 / 24 | [21.63, 127.36] |
| *I. hostmannii* | C0 | D1 | 0.739 | 35 / 32 | [-21.89, 30.71] |
| *I. hostmannii* | C0 | D2 | 0.371 | 32 / 32 | [-14.44, 38.18] |
| *I. hostmannii* | C0 | D3 | 0.056 | 23 / 29 | [-0.78, 63.92] |
| *I. hostmannii* | C0 | R1 | 0.82 | 29 / 27 | [-26.36, 33.17] |
| *I. hostmannii* | C0 | R2 | 0.135 | 26 / 27 | [-9.62, 69.62] |
| *I. hostmannii* | C0 | R3 | 0.478 | 20 / 18 | [-30.75, 64.32] |
| *J. copaia* | C0 | D1 | 0.951 | 34 / 29 | [-49.82, 52.97] |
| *J. copaia* | C0 | D2 | 0.65 | 31 / 23 | [-71.15, 44.8] |
| *J. copaia* | C0 | D3 | **0.022** | 22 / 23 | [15.82, 188.71] |
| *J. copaia* | C0 | R1 | 0.924 | 28 / 24 | [-88.62, 80.6] |
| *J. copaia* | C0 | R2 | 0.725 | 25 / 18 | [-104.09, 73.02] |
| *J. copaia* | C0 | R3 | 0.126 | 19 / 15 | [-23.01, 175.51] |
| *P. officinalis* | C0 | D1 | 0.37 | 35 / 32 | [-27, 71.53] |
| *P. officinalis* | C0 | D2 | **0.036** | 32 / 32 | [4.15, 114.09] |
| *P. officinalis* | C0 | D3 | **0** | 22 / 11 | [147.87, 310.28] |
| *P. officinalis* | C0 | R1 | 0.84 | 29 / 27 | [-56, 68.59] |
| *P. officinalis* | C0 | R2 | **0.012** | 26 / 27 | [20.2, 153.78] |
| *P. officinalis* | C0 | R3 | **0.002** | 19 / 5 | [105.54, 374.37] |
| *S. globulifera* | C0 | D1 | 0.848 | 34 / 27 | [-37.42, 45.36] |
| *S. globulifera* | C0 | D2 | 0.084 | 31 / 30 | [-4.2, 64.62] |
| *S. globulifera* | C0 | D3 | **0.005** | 22 / 26 | [25.65, 132.83] |
| *S. globulifera* | C0 | R1 | 0.607 | 28 / 22 | [-68.68, 40.65] |
| *S. globulifera* | C0 | R2 | 0.997 | 25 / 25 | [-41.46, 41.28] |
| *S. globulifera* | C0 | R3 | **0.001** | 18 / 19 | [46.48, 165.35] |
| *T. melinonii* | C0 | D1 | 0.583 | 34 / 29 | [-14.16, 24.95] |
| *T. melinonii* | C0 | D2 | 0.409 | 31 / 32 | [-33.78, 13.95] |
| *T. melinonii* | C0 | D3 | 0.093 | 22 / 21 | [-5.67, 70.05] |
| *T. melinonii* | C0 | R1 | 0.679 | 28 / 24 | [-26.84, 40.87] |
| *T. melinonii* | C0 | R2 | 0.996 | 25 / 27 | [-33.63, 33.48] |
| *T. melinonii* | C0 | R3 | **0.018** | 19 / 8 | [19.47, 188.29] |
| *V. surinamensis* | C0 | D1 | **0.014** | 35 / 32 | [11.88, 100.27] |
| *V. surinamensis* | C0 | D2 | **0.001** | 32 / 32 | [37.88, 129.57] |
| *V. surinamensis* | C0 | D3 | **0** | 23 / 6 | [320.29, 443.21] |
| *V. surinamensis* | C0 | R1 | 0.422 | 29 / 27 | [-35.93, 84.59] |
| *V. surinamensis* | C0 | R2 | **0.004** | 26 / 26 | [36.01, 177.99] |

**Table S8**: Significance tests for the effects of drought and recovery periods on traits measured on individual plants. Results are from analyses of variance with linear models with *treatment, species,* the interaction *treatment x species* and the plant position *block.sblock* in the greenhouse. Bold p-values indicate significant differences ($p < 0.05$); a p-value of 0 indicates $p < 0.00$.

| Period | Trait | Treatment | pvalue | Species | pvalue | Treatment:Species | pvalue | Block.Sblock | pvalue |
|---|---|---|---|---|---|---|---|---|---|
| Drought | $A_{sat}$ | **30.55** | **0** | **5.32** | **0** | **1.96** | **0.014** | 0.72 | 0.8 |
| Recovery | $A_{sat}$ | 2.54 | 0.06 | **4.29** | **0.001** | **2.22** | **0.008** | 0.92 | 0.5 |
| Drought | $g_s$ | **46.22** | **0** | **5.23** | **0** | 1.19 | 0.3 | 1.05 | 0.4 |
| Recovery | $g_s$ | **5.33** | **0.002** | **5.02** | **0** | **2.96** | **0** | 0.94 | 0.5 |
| Drought | E | **28.44** | **0** | **4.03** | **0.001** | 0.98 | 0.5 | 0.78 | 0.7 |
| Recovery | E | **6.78** | **0** | 1.7 | 0.1 | 1.51 | 0.1 | 1.27 | 0.2 |
| Drought | WUE | **20.96** | **0** | 1.74 | 0.1 | **2.31** | **0.003** | 0.98 | 0.5 |
| Recovery | WUE | 1.1 | 0.4 | **6.01** | **0** | 1.17 | 0.3 | 0.97 | 0.5 |
| Drought | RWC | **128.01** | **0** | **11.67** | **0** | **8.96** | **0** | 0.78 | 0.7 |
| Recovery | RWC | 0.8 | 0.5 | **9.57** | **0** | **3.8** | **0** | 1.76 | 0.06 |
| Drought | $\Psi_{midday}$ | **21.92** | **0** | **21.95** | **0** | **1.91** | **0.023** | 1.45 | 0.1 |
| Recovery | $\Psi_{midday}$ | **3.97** | **0.011** | **34.98** | **0** | **2.6** | **0.003** | 0.61 | 0.8 |
| Drought | Chl | **2.7** | **0.047** | **13.11** | **0** | 0.99 | 0.5 | 0.72 | 0.8 |
| Recovery | Chl | **4.29** | **0.007** | **7.76** | **0** | **1.94** | **0.023** | 1.39 | 0.2 |
| Drought | $F_v/F_m$ | **20.17** | **0** | 0.93 | 0.5 | **2.3** | **0.003** | 1.55 | 0.09 |
| Recovery | $F_v/F_m$ | **10.44** | **0** | **4.78** | **0** | **5.8** | **0** | 0.47 | 0.9 |
| Drought | $L_T$ | **15.87** | **0** | **73.33** | **0** | **2.47** | **0.002** | 0.7 | 0.8 |
| Recovery | $L_T$ | 1.04 | 0.4 | **90.9** | **0** | **3.31** | **0** | 0.28 | 1 |
| Drought | LA | 1.27 | 0.3 | **38.66** | **0** | 0.98 | 0.5 | 1.12 | 0.3 |
| Recovery | LA | **9.54** | **0** | **24.89** | **0** | 0.79 | 0.7 | 0.61 | 0.8 |
| Drought | SLA | 0.5 | 0.7 | **26.52** | **0** | **1.83** | **0.026** | 1.15 | 0.3 |
| Recovery | SLA | **3.29** | **0.023** | **25.37** | **0** | **2.81** | **0.001** | 0.51 | 0.9 |
| Drought | $LA_T$ | **4.2** | **0.007** | **47.4** | **0** | 0.75 | 0.8 | 0.88 | 0.6 |
| Recovery | $LA_T$ | **33.23** | **0** | **39.36** | **0** | 1.18 | 0.3 | 0.28 | 1 |
| Drought | H | **8.06** | **0** | **199.04** | **0** | 1.56 | 0.07 | 1.18 | 0.3 |
| Recovery | H | **31.19** | **0** | **161.86** | **0** | 1.11 | 0.3 | 0.63 | 0.8 |
| Drought | D | **14.78** | **0** | **120.79** | **0** | **1.9** | **0.015** | 1.36 | 0.2 |
| Recovery | D | **31.82** | **0** | **116.57** | **0** | 1.56 | 0.08 | 1.08 | 0.4 |
| Drought | R:S | 2.42 | 0.07 | **16.21** | **0** | **2.26** | **0.004** | 0.98 | 0.5 |
| Recovery | R:S | 2.21 | 0.09 | **27.52** | **0** | 1.51 | 0.1 | 0.97 | 0.5 |

**Table S9:** Weighted degree per trait across treatments. More intense red coloring indicates a higher weighted degree. It is defined as the sum of all significant coefficients of correlation of a node. A node with strength 0 is completely isolated.

| Weighted degree per trait across treatments | | | | | | | |
|---|---|---|---|---|---|---|---|
| | **Control** | **Drought** | | | **Recovery** | | |
| **Trait** | **C0** | **D1** | **D2** | **D3** | **R1** | **R2** | **R3** |
| **Physiology** | | | | | | | |
| $A_{sat}$ | 1.52 | 2.61 | 2.75 | 1.76 | 1.68 | 2.40 | 1.91 |
| $g_s$ | 0.77 | 4.20 | 1.70 | 0.00 | 0.79 | 0.00 | 1.89 |
| *E* | 1.53 | 2.70 | 4.27 | 0.95 | 0.00 | 0.00 | 0.98 |
| *WUE* | 4.84 | 0.81 | 2.59 | 2.55 | 0.90 | 4.23 | 0.96 |
| RWC | 1.54 | 0.79 | 0.86 | 0.76 | 0.00 | 0.78 | 0.00 |
| Chl | 2.36 | 1.67 | 0.86 | 0.00 | 0.00 | 4.26 | 0.00 |
| $F_v/F_m$ | 1.62 | 0.00 | 0.78 | 1.71 | 0.89 | 0.00 | 0.00 |
| **Morphology** | | | | | | | |
| LT | 0.78 | 1.62 | 0.00 | 0.00 | 1.78 | 0.78 | 0.96 |
| LA | 2.48 | 1.57 | 0.83 | 2.38 | 0.78 | 0.00 | 0.00 |
| SLA | 0.76 | 0.78 | 0.00 | 0.00 | 1.63 | 0.00 | 0.88 |
| $LA_T$ | 4.27 | 4.02 | 2.57 | 2.33 | 0.76 | 3.45 | 0.00 |
| H | 3.50 | 2.58 | 1.77 | 3.41 | 1.60 | 3.55 | 2.75 |
| D | 5.16 | 4.97 | 1.81 | 0.91 | 1.74 | 3.49 | 2.80 |
| R:S | 0.78 | 0.84 | 0.76 | 0.00 | 0.93 | 0.00 | 0.00 |
| **Microbiota** | | | | | | | |
| $Div_B$ | 0.82 | 4.05 | 0.83 | 0.00 | 0.00 | 0.00 | 1.84 |
| $Div_F$ | 1.53 | 2.35 | 0.00 | 1.58 | 0.84 | 0.00 | 0.00 |
| $Disp_B$ | 2.62 | 0.00 | 0.76 | 0.00 | 0.84 | 2.55 | 2.84 |
| $Disp_F$ | 4.16 | 2.38 | 0.86 | 1.56 | 0.00 | 1.65 | 2.85 |

**Table S10:** Betweenness per trait across treatments. More intense red coloring indicates a higher centrality. It is defined as the number of shortest paths going through a node. The metric was weighted and normalized to allow for comparison between treatments.

Betweeness per trait across treatments

| | Control | Drought | | | Recovery | | |
|---|---|---|---|---|---|---|---|
| **Trait** | **C0** | **D1** | **D2** | **D3** | **R1** | **R2** | **R3** |
| **Physiology** | | | | | | | |
| $A_{sat}$ | 0.19 | 0.08 | 0.00 | 0.07 | 0.01 | 0.03 | 0.01 |
| $g_s$ | 0.00 | 0.16 | 0.00 | 0.00 | 0.00 | 0.00 | 0.01 |
| *E* | 0.10 | 0.00 | 0.15 | 0.00 | 0.00 | 0.00 | 0.00 |
| *WUE* | 0.24 | 0.00 | 0.05 | 0.18 | 0.00 | 0.04 | 0.00 |
| RWC | 0.10 | 0.00 | 0.00 | 0.00 | 0.00 | 0.00 | 0.00 |
| Chl | 0.26 | 0.00 | 0.00 | 0.00 | 0.00 | 0.03 | 0.00 |
| $F_v/F_m$ | 0.19 | 0.00 | 0.00 | 0.00 | 0.00 | 0.00 | 0.00 |
| **Morphology** | | | | | | | |
| LT | 0.00 | 0.01 | 0.00 | 0.00 | 0.01 | 0.00 | 0.00 |
| LA | 0.10 | 0.01 | 0.00 | 0.17 | 0.00 | 0.00 | 0.00 |
| SLA | 0.00 | 0.00 | 0.00 | 0.00 | 0.01 | 0.00 | 0.00 |
| $LA_T$ | 0.03 | 0.16 | 0.09 | 0.12 | 0.00 | 0.00 | 0.00 |
| H | 0.26 | 0.02 | 0.00 | 0.20 | 0.01 | 0.00 | 0.00 |
| D | 0.07 | 0.15 | 0.00 | 0.00 | 0.01 | 0.00 | 0.00 |
| R:S | 0.00 | 0.00 | 0.00 | 0.00 | 0.00 | 0.00 | 0.00 |
| **Microbiota** | | | | | | | |
| $Div_B$ | 0.00 | 0.05 | 0.00 | 0.00 | 0.00 | 0.00 | 0.01 |
| $Div_F$ | 0.00 | 0.05 | 0.00 | 0.00 | 0.00 | 0.00 | 0.00 |
| $Disp_B$ | 0.00 | 0.00 | 0.00 | 0.00 | 0.00 | 0.01 | 0.00 |
| $Disp_F$ | 0.35 | 0.00 | 0.00 | 0.00 | 0.00 | 0.00 | 0.00 |
